# Evaluating the efficacy of a consumer-centric method for ecological sampling: Using bonobo (*Pan paniscus*) feeding patterns as an instrument for tropical forest characterization

**DOI:** 10.1101/2021.08.04.455026

**Authors:** Erin G. Wessling, Liran Samuni, Roger Mundry, Miguel Adan Pascual, Stefano Lucchesi, Bienfait Kambale, Martin Surbeck

## Abstract

1. Characteristics of food availability and distribution are a key component of a species ecology. Objective measurement of food resources, such as vegetation plot sampling, do not consider aspects of selection by the consumer and therefore may produce imprecise measures of availability. Further, in most animal ecology research, traditional ecological surveying often is time-intensive and supplementary to ongoing behavioral observation. We propose a method to integrate ecological sampling of an animal’s environment into existing behavioral data collection systems by using the consumer as the surveyor. Here, we introduce the consumer-centric method (CCM) of assessing resource availability for its ability to measure food resource abundance, distribution, and dispersion. This method catalogues feeding locations observed during behavioral observation and uses aggregated data to characterize these ecological metrics.
2. We evaluated the CCM relative to traditional vegetation plot surveying using accumulated feeding locations across three years visited by a tropical frugivore, the bonobo (*Pan paniscus*), and compared it with data derived from over 200 vegetation plots across their 50km^2^+ home range.
3. We demonstrate that food species abundance estimates derived from the CCM are comparable to those derived from traditional vegetation plot sampling after approximately 600 observation days or 60 spatially explicit feeding locations. The agreement between the methods further improved when accounting for aspects of consumer selectivity in objective vegetation plot sampling (e.g., size minima). Estimates of density from CCM correlated with plot-derived estimates and were relatively insensitive to home range inclusion and other species characteristics, but were sensitive to sampling frequency (e.g., consumption frequency). Agreement between the methods in relative distribution of resources performed better across species than expected by chance, although measures of dispersion correlated poorly.
4. We demonstrate that while providing a robust measure to quantify local food availability, the CCM has an advantage over traditional sampling methods as it incorporates sampling biases relevant to the consumer. Therefore, as this method can be incorporated into existing observational data collection and does not require additional ecological surveying, it serves as a promising method for behavioral ecological data collection for animal species who re-use space and consume immobile food items.

## 1. Introduction

The abundance, dispersion, and distribution of food resources not only determines species distribution but also has a strong impact on many aspects of an animal’s life-history, physiology, and sociality (e.g., Anholt and Werner 1995; Chapman et al. 2015; Davies and Deviche 2014; Hutto 1990; Lambert and Rothman 2005; Rogers 1987; van Schaik et al. 1993; Vogel and Janson 2007). Due to the core importance of food to an organism, the quantification of food availability and distribution are key considerations across studies and disciplines. Methods used to estimate food resource abundance, distribution, and dispersion are just as varied as the questions which necessitate these quantifications (Szigeti et al. 2016).

Measurement of food resource ***abundance** or **density*** (i.e., estimation of the amount of a resource available in a landscape) depends heavily on the type of resource and scale of interest (Bowering et al. 2018; Morrison 2016). Large scale analyses of abundance typically rely on remotely derived proxies via satellite imagery, but for questions related more immediately to the individual or social group scale, direct measurement of exploitable resources offer more direct insights into the resources available to a consumer (e.g., Foerster et al. 2016; Wessling et al. 2020). While mobile resources may be measured via consumer behavior (e.g., attack rates: Hutto 1990), for static food resources like plants, *abundance* is commonly estimated by sampling subsets of the area of interest. Example methods include transects or vegetation plots/quadrats (Baraloto et al. 2013; Ståhl et al. 2017; Vogel and Janson 2007), with the latter being the most common sampling method in studies of frugivorous or folivorous animals. To then estimate ***distribution***, that is, a calculation of relative resource abundance or density across space within a landscape, sampling may be further stratified across a given area relevant from individuals to populations (e.g., home range, landscape, or region).

Measures of ***dispersion*** (i.e., patterns of clustering or patchiness), such as Morisita’s index (Morisita 1962), are used to quantify the clustering of resources over space within a landscape (Krebs 1999; Stephens and Krebs 1986), often used in the contexts of understanding resource competition and socio-ecological behavior (e.g., Vogel and Janson 2011). Quantifications of food species dispersion are perhaps even more varied in practice and sensitive to the scale relevant to the consumer (e.g., Myers 1978; Vogel and Janson 2011). Dispersion metrics may also require distinct sampling methods tailored to specific questions (e.g., Vogel and Janson 2007), thus potentially requiring supplementary surveying effort to food abundance surveying.

Despite its centrality to animal ecological research, ecological sampling design frequently does not conform to recommended standards nor is adequately validated by animal ecologists (Mortelliti et al., 2010; Szigeti et al. 2016). For example, sampling effort can substantially impact measures of resource abundance, but it is rarely validated whether efforts are sufficient to adequately measure the intended metrics. Further, ecological data collection often requires research effort additional to ongoing behavioral observations and is time intensive and thus infrequently conducted. Snapshots of abundance derived from these efforts in a landscape may therefore be used even many years after they have been collected or may fail to account for temporal variation.

The problem of insufficient quantifications of resource availability may also extend to sampling design. While traditional sampling methods in animal ecology may offer an objective measure of the resources potentially accessible to a consumer, these methods are by design blind to aspects of resource selection by the consumer. Given these disadvantages, the question arises whether there is a way to conduct ecological sampling that is time-efficient within existing behavioral data collection systems and also integrates resource selection criteria of the consumer? Behavioral observation has been used previously as a measure of food availability (Lovette and Holmes 1995; Hutto 1990), dispersion (Vogel and Janson 2011), and preference (Forester et al. 2009), however these methods are either limited in application or still necessitate ecological data collection.

We therefore introduce a consumer-centric method (CCM) for animal behavioral ecology studies which uses the consumer as the survey vehicle to potentially quantify food resources in a landscape. With this method, researchers catalogue food resource locations as they are consumed during the process of behavioral observation. Here, we evaluate the CCM relative to traditional habitat plot data collection using accumulated feeding locations from two social groups of a tropical frugivore, the bonobo (*Pan paniscus*), as a case study. Specifically, we investigated whether behavioral data on feeding locations (trees and lianas) provide a reliable dataset allowing inference about food species’ (1) densities, (2) distribution and (3) dispersion. We additionally assess (4) the minimum sampling effort required and (5) for what characteristics of a food species this method can be considered most reliable.

## 2. Methods

### 2.1 Study Species and Behavioral Observation

Data were collected at the Kokolopori Bonobo Reserve (Fig. 1) on two social groups of bonobos (Ekalakala: EKK, Kokoalongo: KKL) between May 2016 and December 2019. Groups were followed daily for behavioral data collection, during which we collected group feeding locations using a GPS (Garmin GPSMAP 62), and circumference at breast height (synonymous with and hereafter referred to as DBH) of feeding trees ≥20cm diameter and lianas ≥5cm DBH (SI 2.1, 2.2). Due to GPS measurement error and consequently an inability to distinguish individual trees on a small scale, we summarized feeding tree locations of each group into presence or absence of each species in 50 × 50 m cells. We used location data collected with the GPS tracklog function to calculate the home range of both bonobo groups using kernel density estimates (see SI 2.1). These groups share overlapping areas of their home ranges, including 64% and 66% of the home ranges of EKK and KKL, respectively (Samuni et al. 2020). We evaluated whether feeding location datasets were sufficiently sampled and stable by considering accumulation patterns of data per species over time (SI 2.3).

**Fig. 1.**
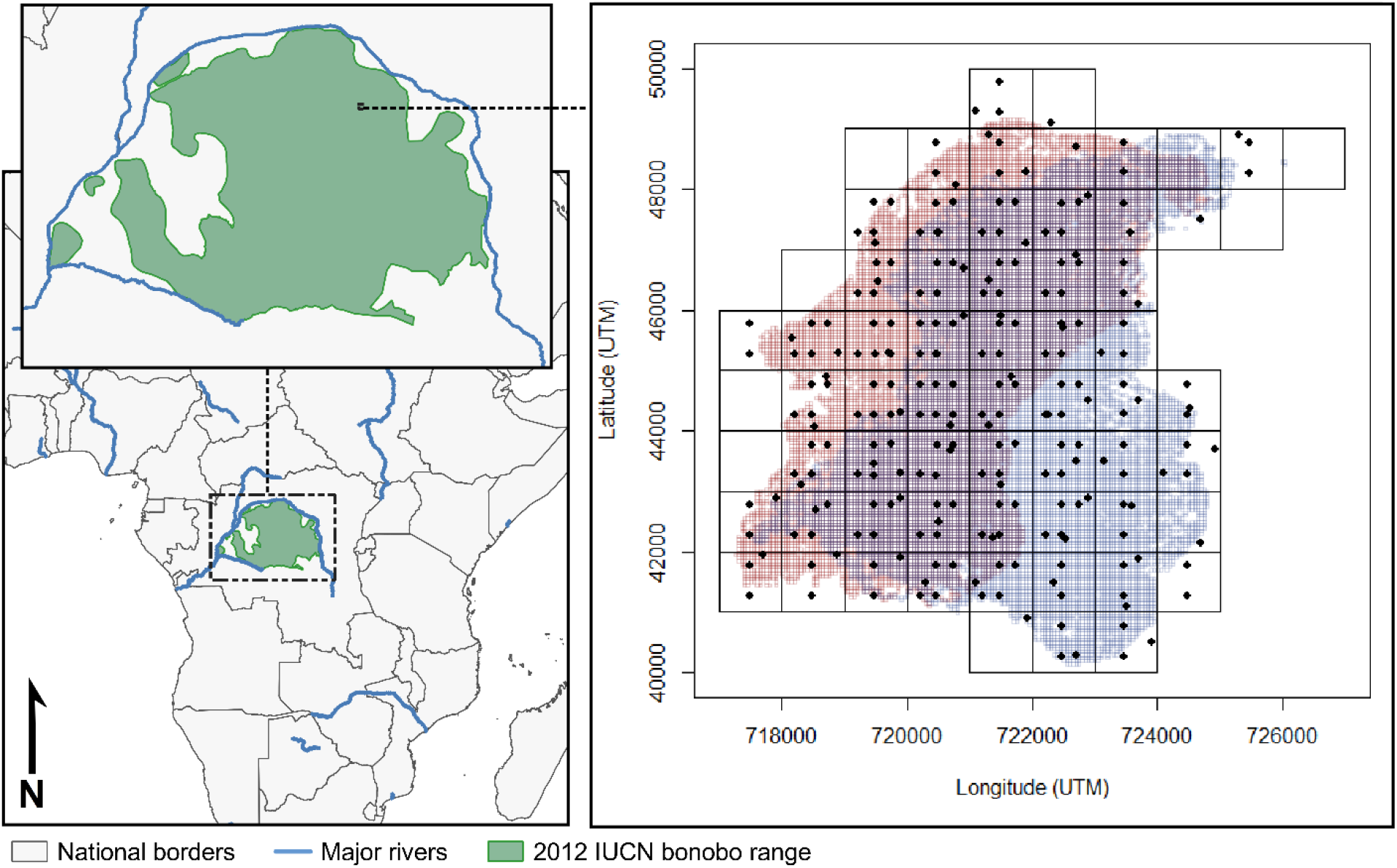
(Left) Location of the study site relative to global bonobo distribution. (Right) 50 × 50m habitat plots (black dots; not to scale) within 1 km^2^ grid cells (black square) overlaid upon all visited 50×50 cells within the 95% home range kernels for Ekalakala (red) and Kokoalongo (blue) bonobo groups.

### 2.2 Vegetation plots

We conducted vegetation plot sampling by overlaying 1×1 km grid cells over the whole ranging area and aimed to conduct plot sampling in every grid cell utilized by at least one of the groups (Fig. 1; SI 1.2). Like the observational cells, all habitat plots were 50 × 50m in size, within which data were collected on all trees meeting the minima defined for observational cells. In total, we sampled 236 plots within these grid cells, of which 214 plots fell within the 95% home range of either group, with 162 and 170 within the 95% range of EKK and KKL, respectively (Fig. 1). Plot sampling averaged 4.1 ± 1.6 (SD) plots per km^2^ (range: 1 to 7) and was determined to be of sufficient sampling depth (SI 1.3).

### 2.3 Comparison of datasets

#### 2.3.1 Density

To compare estimated species abundances derived from each dataset (CCM or vegetation plots), we derived three different indices. 1) We used the bonobo observational data to create a ‘presence index’ based on bonobo feeding locations for each food species, estimated as the number of 50×50m cells in which each species was present divided by the total number of cells within the 95% kernel home range of each group (see Fig. S5 for an example). 2) We calculated species density estimations using the vegetation plot data as the total number of individuals observed per area surveyed (num. individuals / km^2^, hereafter “Plot Density”). 3) We calculated the number of 50×50m plots in which each species was present per total number of plots sampled for more direct comparison with the CCM (hereafter “Plot Presence”).

To evaluate method agreement, we created pair-wise sets of comparisons of the three density indices by means of Pearson’s correlation tests and used the correlation coefficient (r) as a measure of strength of agreement between methods. We conducted the pair-wise comparisons while assessing the influence of sampling effort on method agreement by varying levels of home range usage (kernel % range from 20 until 95 % in increments of 1%) and dietary inclusion (top 10 most consumed species until full diet) for each group. We only considered comparisons with at least 10 species in at least 10 vegetation plots. We additionally created a moving window over the kernel home range from 20% to 95% for which to compare methods more directly according to home range location. This window accounts for variation in area coverage by adjusting window radius to impose similarly sized datasets for comparison over the range of % kernel inclusion (i.e., for agreement from home range core to periphery; SI 3.1).

Finally, to identify potential dataset minima required for reliable and stable density indices derived from the CCM, we evaluated the pattern of correlation strength between indices from each method as the dataset grew over time (i.e., day of data collection), and set the minimum as the point from which the correlation coefficient remains relatively stable. We describe *p*-values for these correlations in our summaries below, however as these correlations require independent data and because we evaluated thousands of correlation coefficients per group (n_EKK_=15075 and n_KKL_=12834), we do not draw inference based on *p*-values but instead focus only upon correlation coefficients.

#### 2.3.2 Dispersion

To evaluate agreement between methods in characterizing food species dispersion, we used Morisita’s index (Morisita 1962). To allow for standardized and directly comparable sample units from which to calculate this index for both methods, we aggregated number of individuals per species visited by the bonobos across three different grid cell sizes (500×500m cells, 1000×1000m cells, and 1500×1500m cells), and calculated the average number of individuals for each species in each of these grid cells using the vegetation plot dataset. For both datasets, we then calculated the Morisita’s index using the *dispindmorisita* function of the package ‘vegan’ (Oksanen et al. 2019) for each species. We further accounted for an unusual distribution of Morisitia’s indices deriving from the vegetation plot dataset by transforming the data to allow for a more normal distribution (SI 3.2).

#### 2.3.3 Distribution

To evaluate the efficacy of the CCM to reliably quantify the distribution of food species in a landscape, we aggregated data by grid cell as in our dispersion comparison. We compiled the abundance data for both bonobo and vegetation plot datasets in two ways: by either i) aggregating (CCM) or averaging (plot dataset) the number of individuals per species per grid cell or ii) by marking the presence/absence of a given species per grid cell size. We chose to average rather than aggregate plot data because greater plot sampling in a grid cell will inherently increase species abundances, whereas sampling biases in CCM could be accounted for by controlling for location within each group’s home range (i.e., % kernel home range). We then fitted model sets separately for each cell size and group (six sets of up to 70 species each), using each food species as a dataset and each cell as a datapoint. We used the estimated bonobo feeding data abundance per cell (a measure of distribution) as the response and the plot abundance as the test predictor using zero inflated Poisson models (500×500m grid size) or simple linear models for (1000×1000 and 1500×1500m grid sizes). Within these models, to account for variation in home range utilization by the bonobos we controlled for the % kernel home range of each cell by averaging the % kernel value assigned to each of the vegetation plots used to estimate the species abundance within that cell. We then calculated average Nagelkerke’s R^2^ (500×500m) or r^2^ (1000×1000m and 1500×1500m) for each model set across levels of dietary inclusion (see SI 3.3 for detailed descriptions of the fitted models and model checks).

To also evaluate agreement between methods on simple presence of a species in a cell, we fitted a generalized linear mixed model with binomial error structure (Baayen 2008) for each grid cell size and each social group. The response in this model was the presence or absence of a species in a given cell as predicted by the bonobo observational data (with one datapoint per species per cell), and presence as measured by vegetation plot and % kernel as test predictors. In these (six total) binomial models we included cell ID and species as random effects and included random slopes for presence/absence in the plots and their correlation within the random effect of species (SI 3.3 for details and model checks). As a last validation of distribution agreement, we identified when bonobos missed the presence of a species in a cell that had been identified in habitat plots and calculated a proportion of missed species occurrences out of all cells per species, as well as evaluated potential sources of biases in likelihood to miss a species in a cell (see 2.4).

### 2.4 Identifying sources of bias

If a consumer is selective in which resources it uses within a landscape, then measurements from vegetation plots may not accurately measure the relevant resources to that consumer. To evaluate these potential discrepancies, we compared food tree and liana sizes (strongly tied to variability in food crop production: Chapman et al. 1992; SI 4) between CCM and vegetation plot data as an example of a potential selective characteristic. We then quantified seven characteristics of each species to evaluate how they contribute to rates of data accumulation and agreement between our sampling methods. Specifically, we considered the lifeform (tree or liana), patterns of dispersion, consumed food item (fruit or non-fruit), seasonality of consumption, density in the landscape, DBH variability, and frequency of consumption (SI 4.1) as test predictors in models with the following responses (SI 4.2): (1) the speed at which data accumulate in the CCM dataset (2) a measure of difference between estimates of density between the methods, and (3) likelihood for bonobos to miss the presence of a species in a cell (SI 4.2).

### 2.4 General Analyses

All data analyses were conducted in R (version 4.0.2; R Core Team 2020), and models were fitted using functions of the ‘lme4’ package (1.1.23; Bates et al. 2015). We report *p*-values between 0.05 and 0.1 as a ‘trend’ for all models to ease issues of dichotomization of significance (Stoehr 1999). To avoid issues of multiple testing when identical models were run across responses which varied only in their summary method (e.g., grid cell size) or dataset (e.g., social group), we describe only patterns which are stable and significant or trending across at least half of each model set; full results for all models as well as further description of all methods and model checks can be found in the SI. We used log transformation to help return predictor (e.g., species density, consumption frequency) and response (all density indices) variables to a roughly normal or symmetric distribution when they were right-skewed.

## 3. Results

### 3.1 Consumer Centric Dataset

The bonobo groups visited (i.e., fed in) a total of 12430 (EKK) and 13827 (KKL) 50×50m cells, amounting to an area ‘surveyed’ of 31.1 (EKK) and 34.6km^2^ (KKL). This amounts to 58.6km^2^ total area surveyed, as 46.7% of this area occurred within the home range overlap of both communities. Bonobos from EKK and KKL fed on a total of 78 tree and liana species (88.6% occurring in the diets of both groups) from trees and lianas, of which 96% of feeding occasions could be identified to a local name. These observations amounted to 8818 (EKK) and 9140 (KKL) unique feeding tree/liana locations (50×50m) consisting of 76 (EKK) and 72 (KKL) species, of which 58 (EKK) and 55 (KKL) species were consumed in at least 10 locations. The diets of both groups were strongly skewed towards a few frequently consumed species (SI 2.4). The groups visited a similar number of locations each day, with a mean of 10.0 ± 5.5 (KKL) and 8.9 ± 5.0 (EKK) locations visited. On average, 4.5 ± 2.0 (KKL) and 4.3 ± 1.8 (EKK) species were consumed per day by the bonobos.

Bonobos visited 60% (EKK) and 56% (KKL) of all visited cells within the first year of data collection, with gradual declines in the accumulation of newly visited cells over the 3+ year study period in both groups and a clear approach towards an asymptote for most of the top 30 species (Fig. S2). We found that the speed at which new feeding locations were added to the dataset also decreased across species (i.e., longer accumulation times) with each passing year for both groups, and that much of the observed decrease in new locations visited over time was likely driven by significant gains early within the dataset (Figs. S2, S3; SI 2.3). Data on species more variable in size (DBH) accumulated slower in EKK than species more uniform in size (but no such relationship was found in KKL), and accumulation was also slower in species consumed for their fruits and in more abundant species in the landscape in both groups (SI 4.3, Tables S2 and S3).

### 3.2 Vegetation Plot Dataset

In total, 14855 trees and lianas were measured across 214 habitat plots (SI 1.1), thus exceeding plot surveying minima (124 plots, SI 1.3). Plot surveying required a cumulative total of 146 team days, averaging to 1.7 ± 0.6 (SD) plots completed per team day (range: 1 – 4). Trees comprise the majority (66.9%) of the individuals measured. This dataset averages to 277.7 individual tree and lianas /ha in across the habitat of these two groups, with 196.1 indiv./ha for food species, and 168.2 indiv./ha for potential food trees that meet bonobo size minima (see below) for the EKK and KKL home ranges collectively.

Seventy-five of the 200 taxa identified in the plots were consumed by at least one of the two groups, with 67 of 72 (EKK) and 70 of 75 species (KKL) in the diet occurring in the plots. Like the bonobo diet, the forest is heavily biased towards a few species, with one species accounting for over 10% of the dataset (‘Bofili’, local name for *Scorodophloeus zenkeri*), and the top 10 most common tree species accounting for almost 40% of all trees and lianas (n=6375, 39.2%). Correspondingly, only 16 species account for over 50% of the individuals in the plots, of which 11 occur in the diet of both groups. Species in the bonobo diet accounted for 67% of the total number of trees or lianas observed in the Kokolopori landscape.

### 3.3 Dataset comparison

#### 3.3.1 Consumer selectivity of tree sizes

Trees visited by bonobos were significantly larger on average than trees measured in the plots (EKK: t=−17.71, p<0.001; KKL: t=−20.38, p<0.001), but by only an average of less than 1cm in both groups (Table S1). For 23.1% of consumed species, we found more individuals in the plots that did not reach the minimum size consumed than those who did exceed this minimum threshold. We subsequently restricted all analyses to trees/lianas that met this threshold, consequently reducing the number of individuals included in plot dataset by approximately 18% in both groups (8891 individuals in EKK and 8685 in KKL; SI 4). Reducing the dataset had a measurable effect on the correlation strengths between estimates of density (see below), with an average improvement of 0.04 for comparison (r) of the CCM estimate with the Plot Presence estimates, and 0.07 improvement in correlation coefficient in the comparison of the CCM estimate with Plot Density.

#### 3.3.2 Density

We found that the density estimates from the CCM and vegetation plots were comparable in both groups (Table 1 and Fig. 2). Patterns of correlational strength between the methods stabilized and smoothed from approximately 50% kernel home range inclusion and above, and when approximately a minimum of 15 species was included in the dataset of both groups. Statistical significance of the correlation was reached in both groups when including ca. 20 of the top species or more. Inclusion of less frequently used areas of the home range to the comparison did not appear to considerably affect the strength of agreement between methods but correlation strength decreased with greater number of species included in the comparison (Fig. 2; Table 1). While we did observe that peripheral areas of the home range generally resulted in lower methodological agreement (Fig. S6), bonobo data appeared largely insensitive to inclusion of the outer reaches of the home range in both groups when included alongside more intensively surveyed areas (i.e., the core range).

**Table 1.** Summary of correlation coefficients (r) between density estimates derived from the CCM and vegetation plot sampling for all significant comparisons above 50% kernel home range and of at least 10 species.

| **(a) Kokoalongo** |  |  |  |  |  |  |
| --- | --- | --- | --- | --- | --- | --- |
| | $r_{\text{mean}} \pm \text{SD}$<br>(all combinations) | $r_{\text{range}}$<br>(all combinations) | Number of<br>species with<br>highest $r_{\text{mean}}$<br>($r_{\text{mean}}$)* | Number of<br>species at $r_{\text{max}}$ | % kernel<br>with<br>highest<br>$r_{\text{mean}}$** | % kernel at<br>$r_{\text{max}}$ |
| Plot Presence and<br>Plot Density | $0.96 \pm 0.02$ | 0.82 - 0.99 | - | - | - | - |
| Plot Presence and<br>CCM | $0.48 \pm 0.04$ | 0.31 - 0.55 | 36 species<br>(0.53) | 10 | 94 %<br>(0.50) | 58 |
| Plot Density and<br>CCM | $0.58 \pm 0.04$ | 0.46 - 0.69 | 36 species<br>(0.65) | 71 | 69 %<br>(0.60) | 51 |
| **(b) Ekalakala** |  |  |  |  |  |  |
| | $r_{\text{mean}} \pm \text{SD}$<br>(all combinations) | $r_{\text{range}}$<br>(all combinations) | Number of<br>species with<br>highest $r_{\text{mean}}$<br>($r_{\text{mean}}$)* | Number of<br>species at $r_{\text{max}}$ | % kernel<br>with<br>highest<br>$r_{\text{mean}}$** | % kernel at<br>$r_{\text{max}}$ |
| Plot Presence and<br>Plot Density | $0.97 \pm 0.06$ | 0.88 - 0.99 | - | - | - | - |
| Plot Presence and<br>CCM | $0.52 \pm 0.02$ | 0.31 - 0.63 | 40 species<br>(0.58) | 48 | 51 %<br>(0.56) | 95 |
| Plot Density and<br>CCM | $0.54 \pm 0.03$ | 0.42 - 0.70 | 19 species<br>(0.66) | 26 | 50 %<br>(0.58) | 95 |

\*averaged across all combinations of % home range inclusion per number of species included
\*\*averaged across all combinations of number of species included per % home range inclusion

**Fig. 2.**
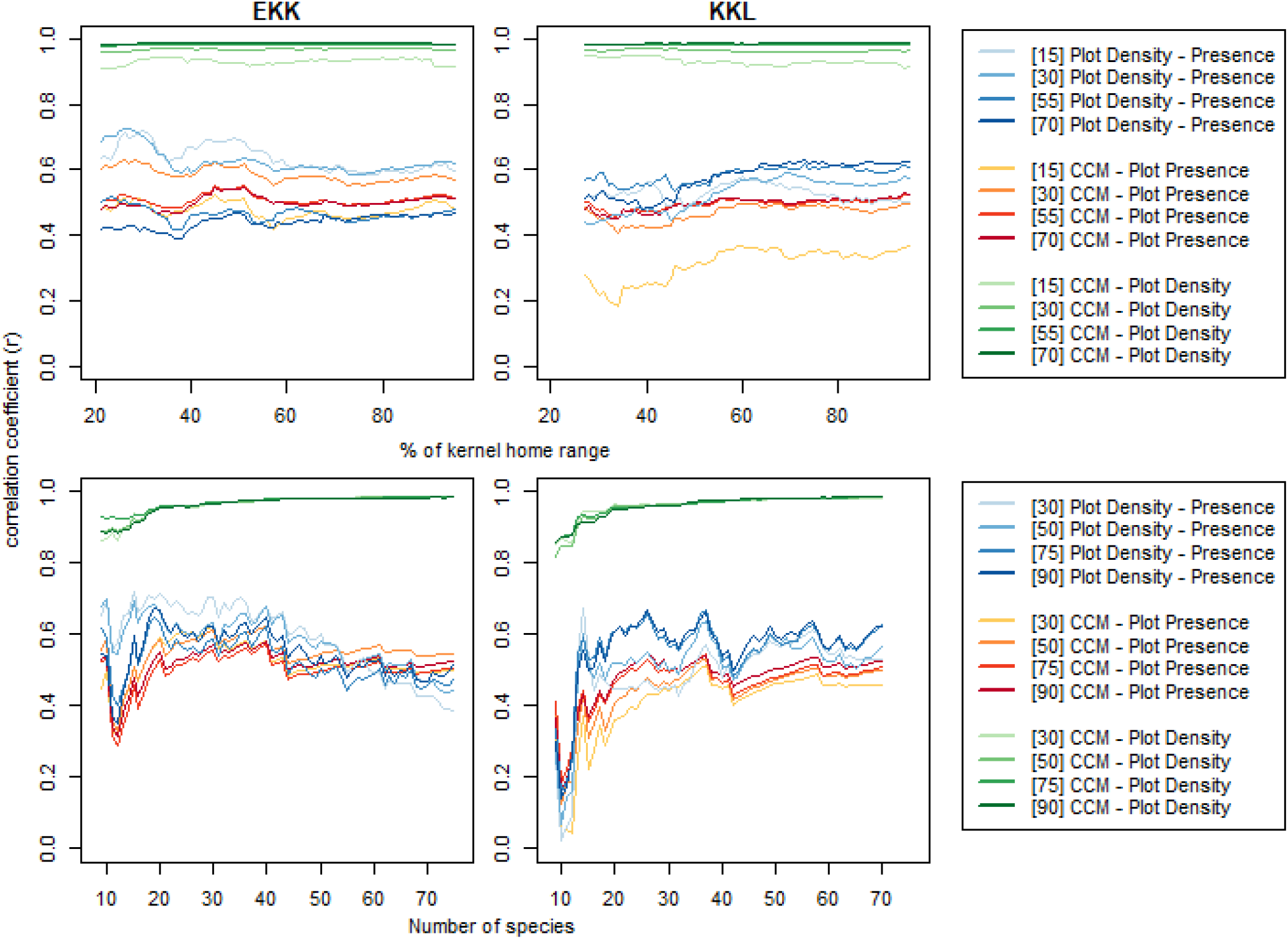
Correlation coefficients of density estimates between sampling methods (i.e., CCM and vegetation plots) for EKK (left) and KKL (right), according to home range percentage (top) and dietary inclusion (bottom). Color groups depict the three comparisons in this study (see legend), with numbers in brackets indicating number of species included (top legend) or percent home range included (bottom legend).

Broadly, the CCM more closely matched estimates of Plot Density relative to Plot Presence. However, for both comparisons we observed a decrease in the correlation coefficient the greater the number of species included in the EKK dataset (Fig. 2, blue lines in bottom left panel). For both groups, we found highest agreement between methods when restricting the comparison to the top 36-40 species (i.e., approximately half of the species in the diet), with one exception that only slightly outcompeted the r of the same range (KKL CCM vs. Plot Density). As expected, comparison between Plot Density and Plot Presence remained consistently high regardless of location within the home range of the bonobo groups, although correlations were lower when fewer species were included.

Once our moving window reached the dataset minimum of 20 plots at ca. 30% kernel, the correlation coefficient of the CCM with plot estimates increased until they reached a maximum around 60% kernel home range in both groups (Fig. S6). Peripheral areas of the home range were generally lower in agreement than more central areas but did not show persistent decreases with increasing peripheralization in a manner that would suggest consistently poorer sampling in peripheral areas. Sampling agreement was strongest within our moving windows for the most frequently consumed species (e.g., 15 or 30 species) relative to more comprehensive subsets of the two groups’ diets (e.g., 55 and 70 species).

The density of the species in the landscape and the variability in size significantly impacted agreement between the methods (Table S4); specifically, lower species density in the plots (estimate average: 0.57 ± 0.11 [SE]) and lower size variability (−1.29 ± 0.62 [SE]) improved method agreement. Further, in KKL only, greater seasonality, non-fruit item consumption, and greater consumption frequency decreased agreement between methods.

Correlation strength between the two methods reached significance and stabilized across methods and groups once exceeding 600 days (i.e., ca. 5300 [KKL] to 6000 [EKK] total visited locations) and continued to improve as data was collected until the end of our data period (Fig. 3; EKK_max_: 1222 days, KKL_max_: 1151 days).

**Fig. 3.**
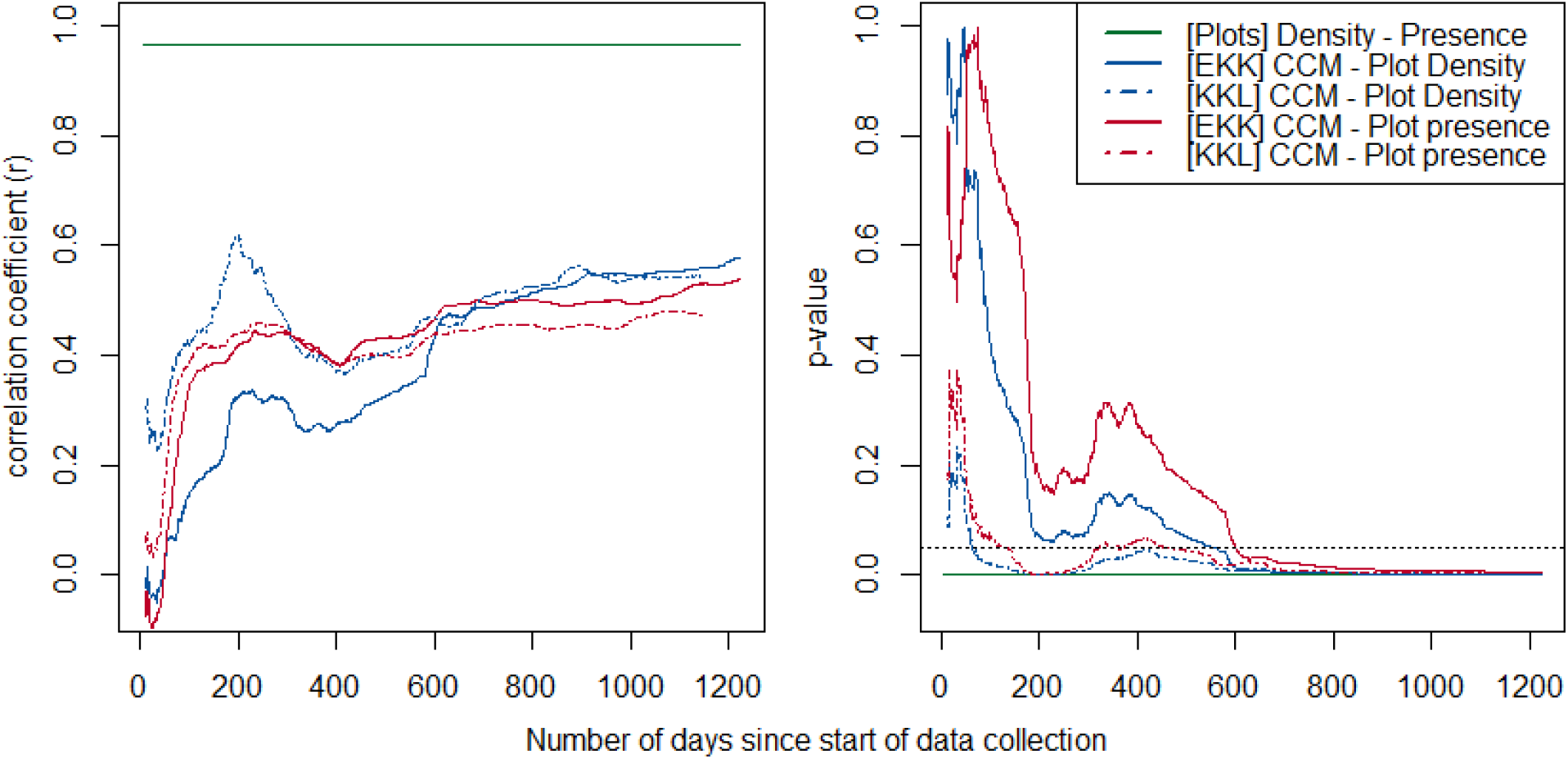
Pearson’s r (left) and p-value (right; dashed line indicates 0.05 alpha level) of all three methods comparisons (see legend) for Ekalakala (full line) and Kokoalongo (dashed lines) over the duration of the dataset.

#### 3.3.3 Dispersion

Overall, Morisita’s indices from the CCM correlated weakly and non-significantly to vegetation plot indices, regardless of grid cell size used or bonobo group (Table 2a).

**Table 2.** Average (a) correlation coefficients (r) and (b) proportion of variance explained (r; 500×500m) or Nagelkerke’s R (1000×1000m and 1500×1500m) between the CCM and plot datasets across three different grid cell sizes for (a) dispersion and (b) distribution estimates.

| (a) Dispersion |  |  |  |  |  |
| --- | --- | --- | --- | --- | --- |
|  | EKALAKALA |  | KOKOALONGO |  |  |
| Cell size | Mean ±SD (range) | | Mean ±SD (range) | | |
| 500 | 0.08 ± 0.17 (-0.54, 0.55) | | 0.00 ± 0.16 (-0.35, 0.61) | | |
| 1000 | 0.00 ± 0.19 (-0.8, 0.25) | | -0.03 ± 0.14 (-0.65, 0.13) | | |
| 1500 | -0.17 ± 0.14 (-0.86, 0.07) | | -0.20 ± 0.14 (-0.83, -0.01) | | |
| (b) Distribution |  |  |  |  |  |
|  | EKALAKALA |  | KOKOALONGO |  |  |
| Cell size | Mean ± SD (range) | Num species $p < 0.05$ (% of total species) | Mean ± SD (range) | Num species $p < 0.05$ (% of total species) | Significant species in both groups |
| 500 | 0.25 ± 0.05 (0.21, 0.38) | 15 (29%) | 0.23 ± 0.04 (0.20, 0.36) | 13 (28%) | 11 |
| 1000 | 0.23 ± 0.02 (0.20, 0.30) | 13 (19%) | 0.24 ± 0.02 (0.20, 0.31) | 8 (11%) | 7 |
| 1500 | 0.23 ± 0.03 (0.14, 0.27) | 8 (12%) | 0.24 ± 0.02 (0.21, 0.27) | 6 (9%) | 3 |

#### 3.3.4 Distribution

Across both bonobo groups and all three grid cell sizes, we found that more species significantly correlated between the two methods for individual abundances across cells than would be expected by chance, with an average of 18% of species significantly correlated between methods across the three cell sizes (Table 2b). Percentage of species with significant correlations across methods declined as grid cell sizes increased, as did the number of significant species which remained consistent across both groups. Generally, proportion variance explained (r or Nagelkerke’s R) by abundance per cell based on plots averaged 0.25 ± 0.32 [SD] across species in all grid cell sizes and groups for predicting abundance per cell based on CCM. Average r did not vary substantially with cell size or between groups (Fig. 5).

**Fig. 5.**
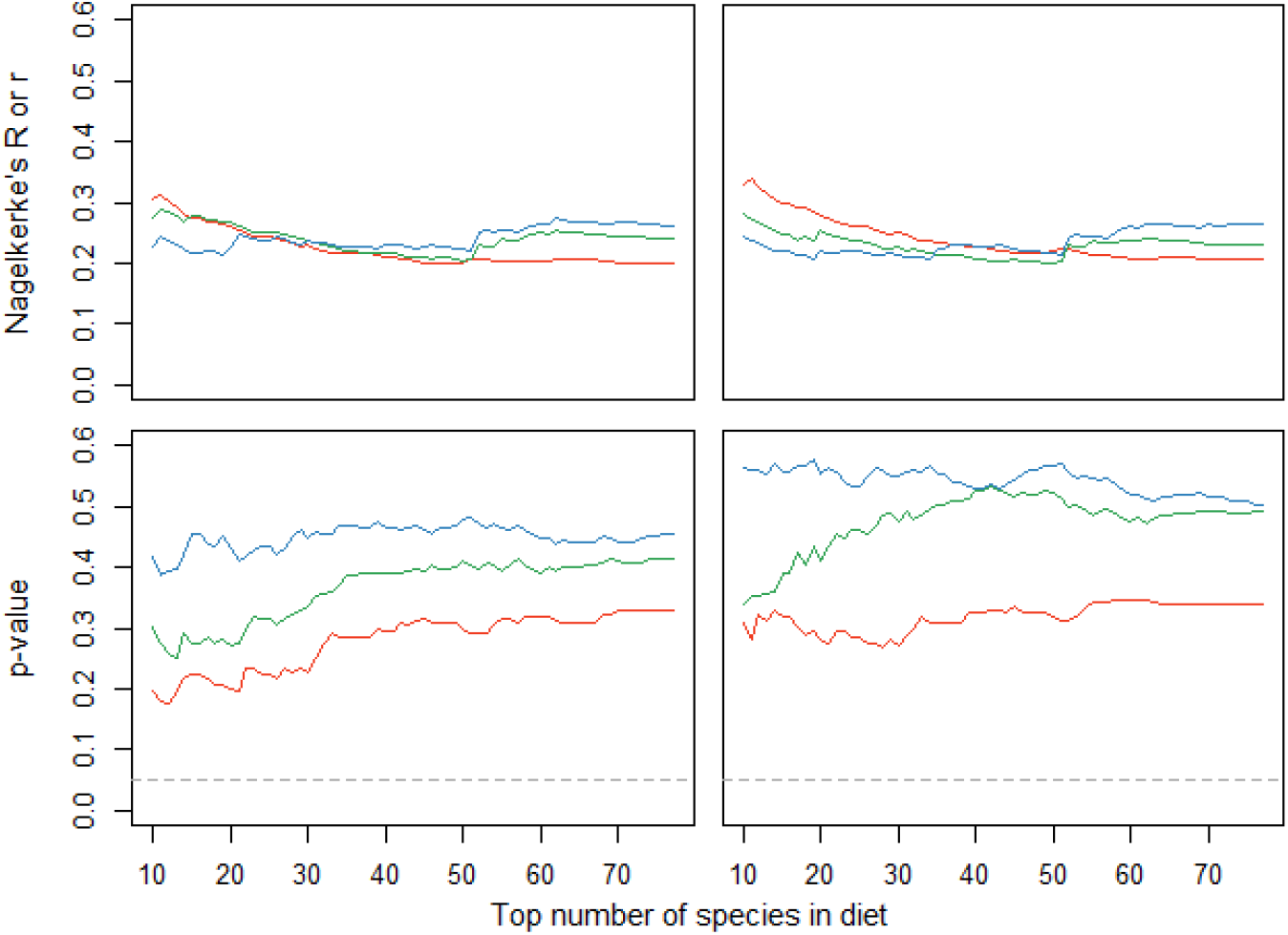
Averaged proportion variance explained (r) or Nagelkerne’s R (top) and p-values for the estimate (bottom; dashed line indicates 0.05 alpha level) for correlations between estimated abundances per cell of species (i.e., distribution agreement) as derived from CCM and vegetation plots for EKK (left) and KKL (right) and for three grid cell sizes (red: 500×500m, green: 1000×1000m, blue: 1500×1500m).

The presence of a species in a cell as measured by plots significantly predicted the presence of that species in the cell as identified with CCM (estimate: 0.60 ± 0.20 (SD), range: 0.32 – 0.81; Table S5). The location of a cell within the home range appeared to play a consistent role, with food species less likely to be identified by CCM in more peripheral cells (average estimate: −0.05 ± 0.01 (SD), range: −0.05 – −0.04; Table S5). Bonobos missed presence of a species on average in 17.5% ± 16.3% (SD; range: 0 – 68.4%) of the 500×500m cells and in 18.4% ± 16.5% (SD; range: 0 – 61.2%) of the 1000×1000m cells. Increases in species abundance correlated with an increase in the likelihood for bonobos to miss the presence of species in a cell irrespective of cell size or group but species were less likely to be missed in a cell if they were more frequently consumed. We additionally found some support for species consumed for their fruits to be more likely to be missed in smaller cell sizes (Table S6).

## 4. Discussion

Here we demonstrate the applicability of the consumer-centric method (CCM) for measuring resource density and distribution in an animal’s landscape. We demonstrate that food species estimates derived from the CCM method are comparable to estimates derived from traditional vegetation plot sampling following a relatively short data collection timeframe, including before data have reached saturation. The method also seems promising for characterizing distribution of food patches within a landscape. Furthermore, we demonstrated that the CCM has an advantage over traditional sampling methods as it incorporates sampling bias important to the consumer into quantification of the ecological landscape.

### 4.1 Robustness of the CCM

The CCM estimates of abundance showed strong similarity to estimates from traditional ecological sampling. Behavioral ecologists have previously used consumption rates to infer about the abundance of food resources (Hutto 1990; Lovette & Holmes 1995; Watts & Mitani 2015). These methods are particularly susceptible to handling time, consumer motivation, and/or dependence of preference from resource availability and are subsequently difficult to validate (Lovette & Holmes 1995). The key advantage of the CCM is that rather than quantifying availability from occurrences of consumption (frequency dependent) it depends on independent locations (spatially dependent), thereby allowing validation with traditional vegetation plot sampling.

Although we found a significant but minor periphery effect on agreement between methods in the presence/absence of species, the correlation of abundance estimates between methods were unaltered by % of home range inclusion. The lack of a spatial effect on agreement between the methods is in some part likely to be a result of home range selection on the part of the consumer (e.g., second-order selection *sensu* Johnson 1980), i.e., bonobos may have already selected their home range based upon resource availability hence no sampling biases therewithin. In absence of home range use biases, the CCM therefore reliably estimates resource availability across the entirety of a group’s space use, although future studies should verify absence of sampling biases on agreement between the CCM and traditional methods in their own study species.

Further, we found consumption frequency to impact likelihood to miss species presences. Consequently, restricting estimation to only the top half of the consumed species (by frequency) appears to offer a compromise between maintenance of dietary relevance while maximizing fidelity with density estimates as assessed by objective plot measurements. This minimum translated to species consumed in approximately at least 60 locations over our three-year dataset. A general consequence of sampling frequency by a consumer is that estimates improve in precision as data accumulate over time. While species in our dataset were variable in “saturation level”, rates of new locations sampled by the bonobos slowed over the course of data collection and inter-method correlation of species abundances stabilized after fewer than two years of data collection (approx. 600 days). As our results indicate that sampling rate affects stability of estimates (e.g., frequency of consumption), we anticipate that this general minimum will be higher for species with slower sampling frequency, i.e., for less frequently consumed species, as well masting species or species which are consumed aseasonally.

Generally, species distribution (i.e., relative abundance) correlated weakly between the methods across species regardless of scale of comparison (i.e., cell size). A greater proportion of species reached significant agreement between methods in smaller rather than larger cell sizes, potentially as a function of proximity, i.e., the larger the cell size used the greater the potential distance between bonobo feeding locations and comparatively small plot areas. Nevertheless, our finding that correlations of distribution within species was significant across a greater proportion of food species than expected by chance (i.e., 5%) and that the rates at which bonobos missed the presence of a species in a cell is likewise better than common rates of species misses between multiple observers in single plot (Millberg et al. 2008) provides hope that reliable estimates of sub-landscape abundances may improve with greater sampling depth.

While detectability is rarely 100% in either method (Morrison 2016), the miss rates by a consumer in the CCM may rather carry additional information about the nature of resource selection (and the individuals which are subsequently ignored). This is especially likely to be the case in consumers who have the capacity to keep track of spatiotemporal patterns of resource availability. Bonobos likely have a concept of where and when resources become available, and therefore are also capable of targeting resources that are rare (Janmaat et al. 2013; Normand et al. 2013). Consequently, the CCM mimics *ad hoc* sampling (Foster et al. 1998, Gordon & Newton 2006, Hopkins 2007), and our results indicate that the CCM more closely matches plot density estimates at capturing rare species relative to more abundant species.

Nonetheless, in absence of full censusing, we cannot differentiate between which sampling method produced a more precise representation of food species availability, dispersion, and distribution patterns. Ideally, methodological sampling biases could be identified by simulating both sampling schemes from a simulated ‘forest’. Unfortunately, we rarely understand the complexity of consumer movement and resource selection patterns (Buskirk & Millspaugh 2006). Therefore, subsequent conclusions drawn from simulated sampling behavior would be just as arbitrary as the decisions made to simulate them (Johnson 1980).

### 4.2 Measuring different phenomena

We argue that the CCM, with adequate evaluation, may be a more appropriate tool for most applications in behavioral ecology than traditional inventory methods. Traditional plot sampling quantifies the total amount of potential resources which also include inaccessible, unattractive, or otherwise unpalatable resources to a consumer. Only a subset of these resources comprises true resource availability, i.e., resources with potential to be selected (Alldredge et al. 1998; Buskirk & Millspaugh 2006; Johnson 1980), and although correlated, each represents inherently two separate phenomena (Hutto 1990). Because we rarely understand the processes of food selection by which consumers filters objective resource abundance into availability, the CCM provides the advantage of using the consumer as a means to avoid arbitrary decisions as to how to best sample the landscape (Johnson 1980). We detail examples of this selectivity and the resulting advantages of the CCM below.

First, we observed significant differences between average sizes of trees/lianas visited by bonobos relative to what was available in the landscape of consumed species (as measured in vegetation plots). Reducing our plot dataset to sizes selected by the consumer increased the correlations between CCM and vegetation plot measures and demonstrates the inadequacies of consumer-objective plots in mirroring consumer behavior. Second, that bonobos missed or ignored certain food resources in cells identified to contain them underlines further how researchers are likely unaware of relevant selection criteria that impact measurement of true resource availability. Because apes possess mental maps of their environments and are known to adjust travel to target preferred food sources (Janmaat et al. 2013; Lucchesi et al. 2021), they are unlikely to consistently miss potentially important food resources within their home range at the scale observed in our study. Third, we found that CMM estimates of density and distribution differed between bonobo social groups, even with largely overlapping home ranges. This conforms to previous findings of group-specific feeding selection criteria in bonobos (Samuni et al. 2020), independent of local abundance. If resource availability for a consumer in a given landscape is dependent on group identity, then only methods like the CCM incorporating these criteria allow comparable estimates for comparative studies across social groups.

Altogether, by accounting for consumer selection, the accumulation of data on food patch location are inherently less subjective than datasets dependent upon arbitrary decisions by the investigator (Johnson 1980). Biases in resource measurement occur via multiple sources including selection of sampling method, metric, effort, as well as through unavoidable systematic or random measurement errors (Baraloto et al. 2013; Milberg et al. 2008; Morrison 2016; Ståhl et al. 2017; Wessling et al. 2020). The CCM, however, accounts for several of these issues because consumers are knowledgeable and motivated surveyors who actively target resources, with apparently negligible impact of scale variation (e.g., cell size) or abundance on fidelity of CCM estimates to plot-derived estimates.

Therefore, estimates derived from the CCM provide accurate measures of availability once data have reached a sufficient depth. Our spatially explicit CCM further allows for data accumulation and consequential improvement of the accuracy of estimates over time until otherwise removed due to irrelevance (e.g., patch loss). Nevertheless, if rapid abundance assessment is preferable for a project, traditional ecological sampling may remain a preferable method due to a 600 person-day burn-in time required (this study) by the CCM before estimates become reliably stable per social group relative to 150 person days of plots for both groups. However, these 150 person days are supplementary to observational data, insomuch as person-days necessary to collect both sets of data must be considered additive to observational data collection. Yet, if databases of feeding locations are already available, adapting these data to CCM estimation of resource density or distribution save researchers from needing to collect additional data to quantify resource abundance.

While this method is best applied to estimate the availability of discrete, immobile, and spatially explicit resources, these advantages transcend application beyond bonobos and allow researchers to evaluate the strengths of the method for their investigations across all potential consumers who meet these criteria (further discussed in Table S7). Functionally, assumptions of the CCM are similar to studies investigating resource preference, a method which also combines objective habitat measures with subjective animal-centric data (Manly et al. 2002). For example, this method can only be applied to consumers which re-use space over time, like a consistent home range, and assumes that consumers have equal access to all the areas of this space (Alldredge et al. 1998). Nonetheless, researchers must verify CCM sampling is of sufficient sampling depth and absent of biases (e.g., sampling biases or characteristics of food items) for their consumer before the CCM can be applied as a means of resource availability. When applied correctly, the CCM will enable many behavioral ecologists to quantify aspects of food availability by using existing data, in a manner that is more suitable to its application as well as allows for more precise comparison ways that make this data comparable across social groups, subsequently promising new insights in the interplay between an animal and its environment.

## Author contributions

EW, LS conceptualized the manuscript, EW, LS, MS designed the analyses, EW, LS, RM performed the analyses; EW, MAP, SL, BK, MS provided data; EW wrote the manuscript with input from all co-authors.

## Statement of where you intend to archive your data

All necessary data to reproduce this study will be submitted to the Dryad Digital Repository.

## Acknowledgements

This work is dedicated to the late Dr. Deborah Moore, who is deeply missed at Kokolopori. We are grateful to the Ministry of Scientific Research and Technology (MSRT) and the ICCN of the Democratic Republic of the Congo, as well as to residents of the surrounding communities for facilitating our research in the Kokolopori Bonobo Reserve. We thank Leonard Nkanga, Paulin Lokutshu Lokole, Serge Betamba Bokelakela, Valerie Beraud, Vincent Rison, and the team of local assistants for their invaluable contribution to data collection and bonobo tracking. This work was supported by Harvard University, and meets ethical standards for animal research of the Max Planck Society and the MSRT. The authors have no conflict of interest to declare.

