## Supplementary Information for "Evaluating the efficacy of a consumer-centric method for ecological sampling: Using bonobo (*Pan paniscus*) feeding patterns as an instrument for tropical forest characterization"

### SUPPORTING INFORMATION

### 1. Vegetation Plot Sampling

#### 1.1 Sampling Structure

We conducted vegetation plot sampling of the Kokolopori Bonobo Reserve (Surbeck et al. 2017) as one of several commonly used methods for sampling arboreal density, dispersion, and distribution in animal research. To do so, we overlaid 1x1 km grid cells over the home ranges of both groups and aimed to conduct plot sampling in every grid cell utilized by at least one of the groups. We sampled a total of 236 plots across the study site for all trees ≥20cm diameter breast height (DBH) and lianas ≥5cm DBH. Tree and liana species were identified by observers using their names in the local language (Longando), and in all subsequent analyses we continued to label species by their Longandan names, which have been taxonomically identified by local botanists. Few Longandan names refer to multiple species belonging to the same genus or (rarely) from separate genera, but individual species are never identified by more than one local name.

16269 trees and lianas were measured across the 236 habitat plots, 99.9% of which were identified to 200 local (Longanda) names (n=21 unidentified). 214 of the 236 plots occurred within the 95% home ranges of the two bonobo study groups (n=14855 trees). Habitat plot data were collected by three different teams, with overlap in membership across two of three teams (i.e., teams of individuals A and B, B and C, and D and E). The three teams collected data using different plot placement methods but identical data collection within the plots. The first dataset (n=65 plots) was collected in 2017, with one plot placed randomly within each grid cell. This dataset was complemented in 2019 by plots systematically placed equidistant down the midline of the 1x1km grid cell (N to S; n=102 plots) and was further supplemented by a third dataset (n=65 plots) which focused on a subset of 30 contiguous grid cells which overlapped with the areas of highest use by both social groups. Plots in the third dataset were placed 250m directly to the west or east of plots from the second dataset, with a goal for a minimum of two plots sampled per each cell and a maximum of four.

1.2 Inter-surveyor reliability

To ensure that data collected by teams of differing membership are comparable and free of bias, we evaluate measuring consistency between two of our vegetation plot sampling teams (i.e., team AB and DE) by comparing data from four identical 50 x 50m plots spread throughout the home ranges of the two social groups. We evaluated agreement between these four plots across several metrics and found that inter-observer reliability of these plots indicated strong but imperfect agreement between the two sets of data collectors. We found that teams differed in the total number of individuals counted in each plot (mean: 83.0 ± 16.5 (SD) individuals/plot; range: 68 - 120) by an average of 12.7 ± 15.7% (SD; range: 0 – 35%), however this mean was inflated largely by a single plot in which liana count was considerably exaggerated by one team, but in which tree count was nearly identical between both teams. Agreement in liana counts (mean: 45.8 ± 5.8 (SD) individuals/plot; range: 37 - 52) appeared to be more variable between observer teams, while average tree count difference between doubly-measured plots averaged only 6.7 ± 4.7% difference (SD; range: 0 – 10%).

Surveying teams also differed in the number of species identified in each plot by an average of 7.6 ± 10.0% (range: 0 – 22%) of species (mean: 31.9 ± 5.0 (SD) species/plot; range: 24 - 39), corresponding to an average of a difference of 2.3 ± 2.6 (SD) species per plot. Species both measured and missed by surveyors within the four plots totaled 17 unique species and did not follow any apparent patterns in bias of species characteristics, abundance, or rates of consumption by bonobos, but the majority of which (n=13) were tree species. Plot averages of tree circumference differed by an average of 3.5 ± 3.4 cm (SD; range: 0 – 6.6cm), but total population differences in tree circumference differed by an average of 8cm between teams (Team 1: 68 ± 71cm, Team 2: 74 ± 66cm).

1.3 Ensuring sufficient sampling effort

To ensure sufficient vegetation plot sampling effort, we defined a threshold after which quantification of landscape variables remained consistent and representative of true landscape-level values. We applied a method we have previously developed (Wessling et al. 2020) to identify a minimum sampling threshold for reliable estimation of three landscape variables: i) tree density, ii) standard deviation of number of trees per plot (as a proxy of landscape heterogeneity), and iii) number of tree species in the dataset (i.e., species accumulation). Succinctly, this method applied simulations of randomized selections of vegetation plots (1,000 iterations, with replacement) across the range of possible dataset sizes (1 to 236 plots), after which we calculated each of the aforementioned landscape variables. For each sampling size we quantified the percentage of values that appear outside of the bounds of the range of the largest dataset (i.e., 236 plots with replacement). We then identified the minimum number of vegetation plots for which at least 95% of randomized selections estimated a value of tree density or SD of number of trees per plot within the range of the largest dataset consistently in every sample size above this threshold. To quantify the accumulation in the number of species identified according to varying number of plots sampled, we used the 'specaccum' function from the package *vegan* (Oksanen et al. 2019) using the 'random' method.

Minimum number of plots required to reach 95% agreement with the maximum plot dataset was 104 plots for consistent variability in number of trees (i.e., SD of trees per plot) and 124 plots for consistent estimates of tree density (Fig. S1). With regard to the number of species identified in plots, we found that species accumulated quickly, necessitating on average a minimum of 167 plots to arrive at 95% of the total possible assemblage identified in our plot data, although arriving already at an estimated 68% of species identified in a 30-plot sample (Fig. S1). The diversity of the species consumed by bonobos accumulated far quicker, with an average of 93 plots required to arrive at 95% total consumed species diversity in the sample, but only 16 plots needed to arrive at 80% species diversity. Given that 114 and 116 plots were sampled within the 95% kernel home ranges of the Ekalakala bonobo group (EKK) and the Kokoalongo bonobo group (KKL), respectively, we can be confident that plot data from within the home ranges of each group are of reasonably sufficient size to provide reliable estimates of abundance and clearly sufficient size to estimate consumed species diversity in each range.

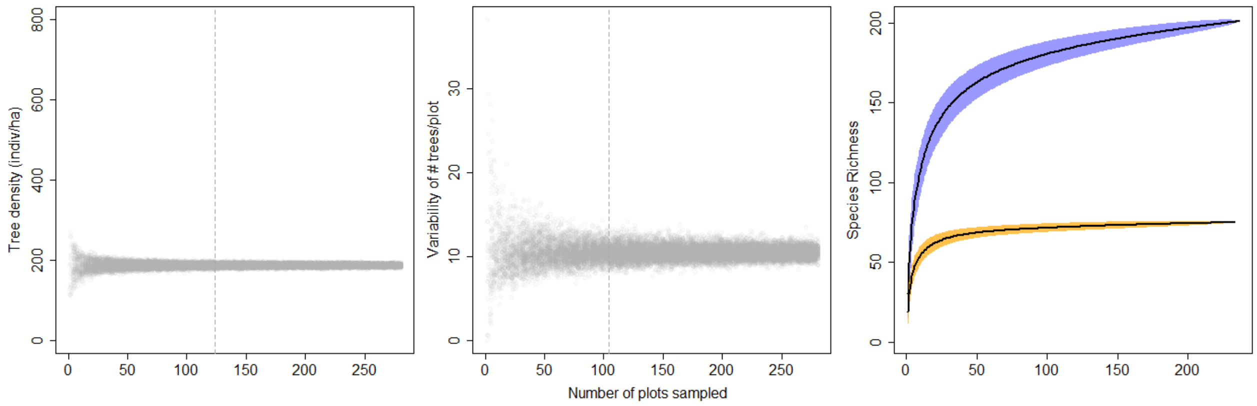

***Fig. S1***. *(Estimates of 1000 iterations per number of vegetation plots of two environmental variables (tree density, left; SD of number of trees per plot, middle) across the Kokolopori Bonobo Reserve. Dotted vertical lines indicate minimum threshold for estimates reaching 95% of the precision of full dataset estimates. The right subplot depicts species accumulation curves for all tree and liana species (blue confidence intervals) and food species only (orange confidence intervals) over number of vegetation plots, with solid black lines representing sampling averages.*

**2. Bonobo Observational Sampling (The CCM)**

2.1 Observational Data

Both groups were followed daily for behavioral data collection for the duration of their full day, during which we collected location data at 1-minute intervals using a handheld GPS device (Garmin GPSMAP 62). During full-day follows, we recorded all fruit tree and liana species fed upon by the bonobos. We additionally collected data on the GPS location (UTM, zone 34N) and circumference at breast height (1.3m; synonymous with and hereafter referred to as DBH) of feeding trees (≥20cm in diameter) and lianas (≥5cm diameter). We recorded the GPS location for 78% (EKK) and 62% (KKL) of the feeding trees and lianas identified, and the DBH in 45% (EKK) and 44% (KKL) of feeding locations. If coordinates of a given tree/liana were not recorded, we obtained the coordinates of that feeding location from time-matched locations of 1-min GPS tracklog data continuously collected. The mean (± SD) time difference between feeding observations and GPS tracklog locations was 45 ± 58 seconds (EKK) and 52 ± 52 seconds (KKL). We verified that we could reliably use the GPS tracklog data as the GPS coordinates of feeding locations by using the dataset of feeding trees and lianas with known locations (GPS data recorded) and calculating the spatial distance between the known waypoint and tracklog locations. The mean (± SD) distance between the recorded tree/liana and tracklog GPS locations was 10 ± 11 meters in Ekalakala and 10 ± 12 meters in Kokoalongo, well within the range of the 50x50 m cells.

We used the bonobo tracklog data to determine the home range distributions of both bonobo groups using kernel density estimates (Worton 1989) produced with the function *kernelUD* of the package ‘adehabitatHR’ (Calenge 2006). The home range sizes (95% kernel) of the two groups during our study period were 35 $km^2$ (EKK), and 40 $km^2$ (KKL). These two social groups share overlapping areas of their home ranges, including 64% (EKK) and 66% (KKL) of group home ranges.

2.3 Ensuring sufficient sampling effort

To assess whether our feeding location datasets were sufficiently sampled and stable, we evaluated for each species consumed by each bonobo group the accumulation of new locations (i.e., 50x50m cells) of that species over the duration of sampling. We considered that a decrease in number of new cells added over the duration of sampling would be indicative of dataset saturation. We quantified this for each species by calculating the proportion of the number of new cells gained within each year since the first occasion of consumption by bonobos in the dataset. We then calculated the slope of the change in this proportion over the three or more years of data collection using a spearman correlation test. If accumulation generally slows, the slope of the proportion gained each year will be negative. We considered the slope for only those species which were consumed in at least 10 locations in subsequent analyses.

We found that the speed at which new feeding locations were added to the dataset (i.e., accumulation slope) decreased consistently across species with each passing year for both groups, and that much of the observed decrease in new locations over time was likely driven by significant gains early within the dataset (Figs. S2, S3). For species which were visited on at least 10 occasions, the change of the slope of new locations identified each year against sampling date was negative across the vast majority of species in both EKK (52 of 55 species, mean change of slope: -0.7, range: -1 to 1) and KKL (51 of 56 species, mean change of slope: -0.6, range: -1.0 to 0.8), although none yielded significant correlations of the slope with sampling year, as the limited sample size allowed only a minimum *p*-value of 0.08 ($n_{max}$=4 calendar years) and was calculated using an exact test (thereby eliminating tests with tied datapoints). Regardless of this overall pattern, feeding tree locations generally did not form a visual asymptote for many species in the bonobos diet, but this pattern was variable across species (see Figs. S2, S3). Some species (e.g., '*Liloma'*, the most frequently consumed arboreal species in the EKK diet) showed clear drop-offs in new locations added to the dataset as time progressed, however a significant proportion of new locations were visited towards the end of our dataset in others (e.g. '*Bolingo'*, the most frequently consumed species in the KKL diet). In some species, bonobos continued to visit many new locations even after three years of data collection (Fig. S2, S3).

Additionally, accumulation of new locations for a species over time could derive both from the visiting of new areas or by repeated surveying of already visited cells. To therefore differentiate between these possibilities, we evaluated the pattern of new cell accumulation in the dataset over time. We found that bonobos visited 60% (EKK) and 56% (KKL) of all visited cells within the first year of data collection, with gradual declines in the accumulation of newly visited cells over the 3+ year dataset in both groups and a clear approach towards an asymptotic pattern indicating sufficient sampling effort of the area at large (Fig. S4).

### 2.4 Bonobo dietary skew

Most feeding locations occurred within the 95% home range kernels of both groups (EKK: 95.6%, KKL: 96.4%). The diets of the two groups were similar, with 70 species (88.6%) occurring in the diets of both groups, and 13 of 15 of the most frequently consumed species dominating the diets of both groups. The diets of both groups were considerably skewed with the top five tree and liana species in each diet comprising 44% (EKK) and 46% (KKL) of feeding bouts on trees and lianas. This skew extended further, to 76% (EKK) and 67% (KKL) of feeding bouts when considering only the top 15 species.

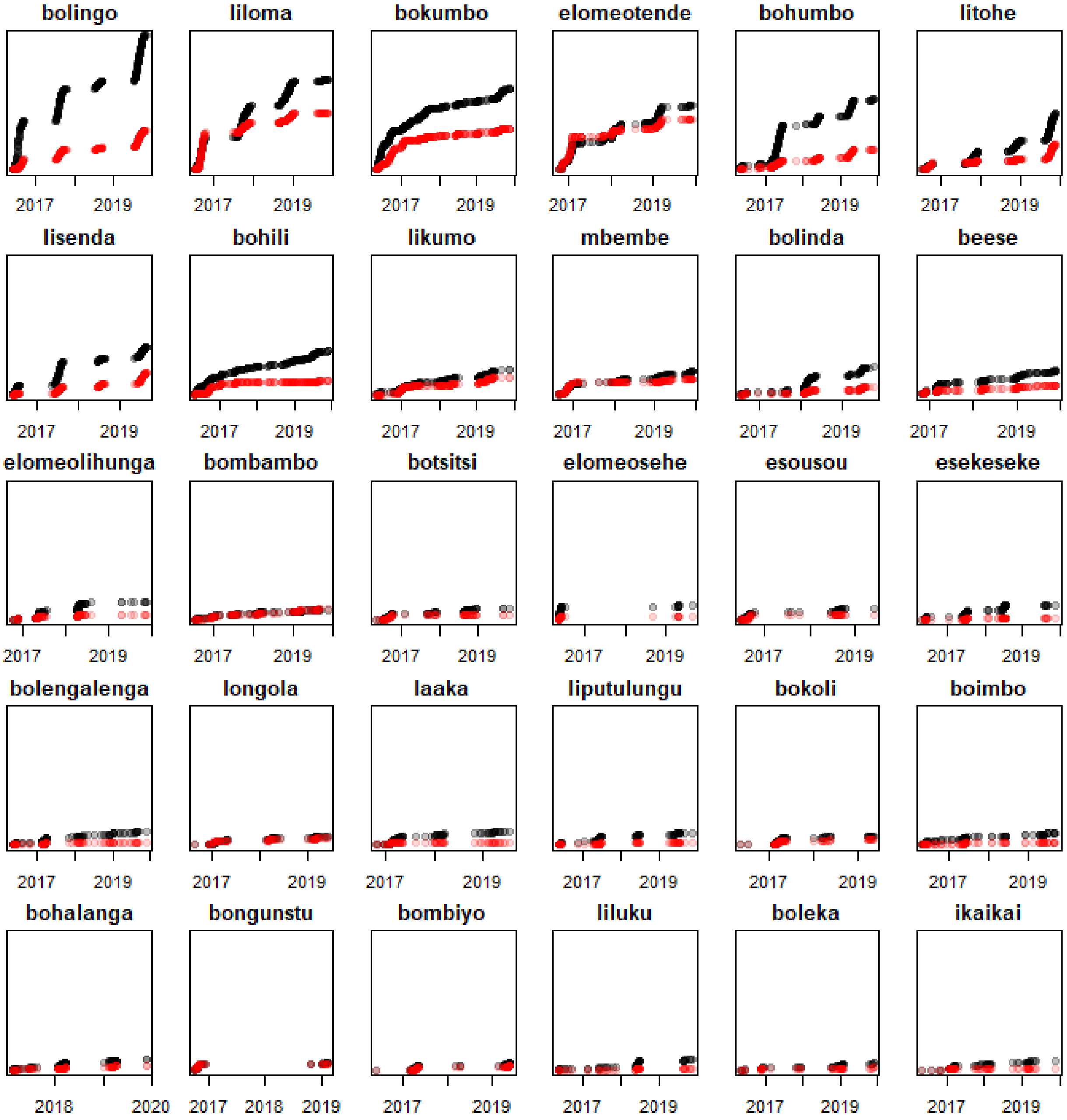

***Fig. S2.*** *Accumulation of new cells containing each species over time in the dataset (black) and accumulation of total number of revisits (red) for the top 30 species in the Kokoalongo (KKL) diet. Y-axis depicts an absolute scale of number of locations visited (range: 0 to 1250).*

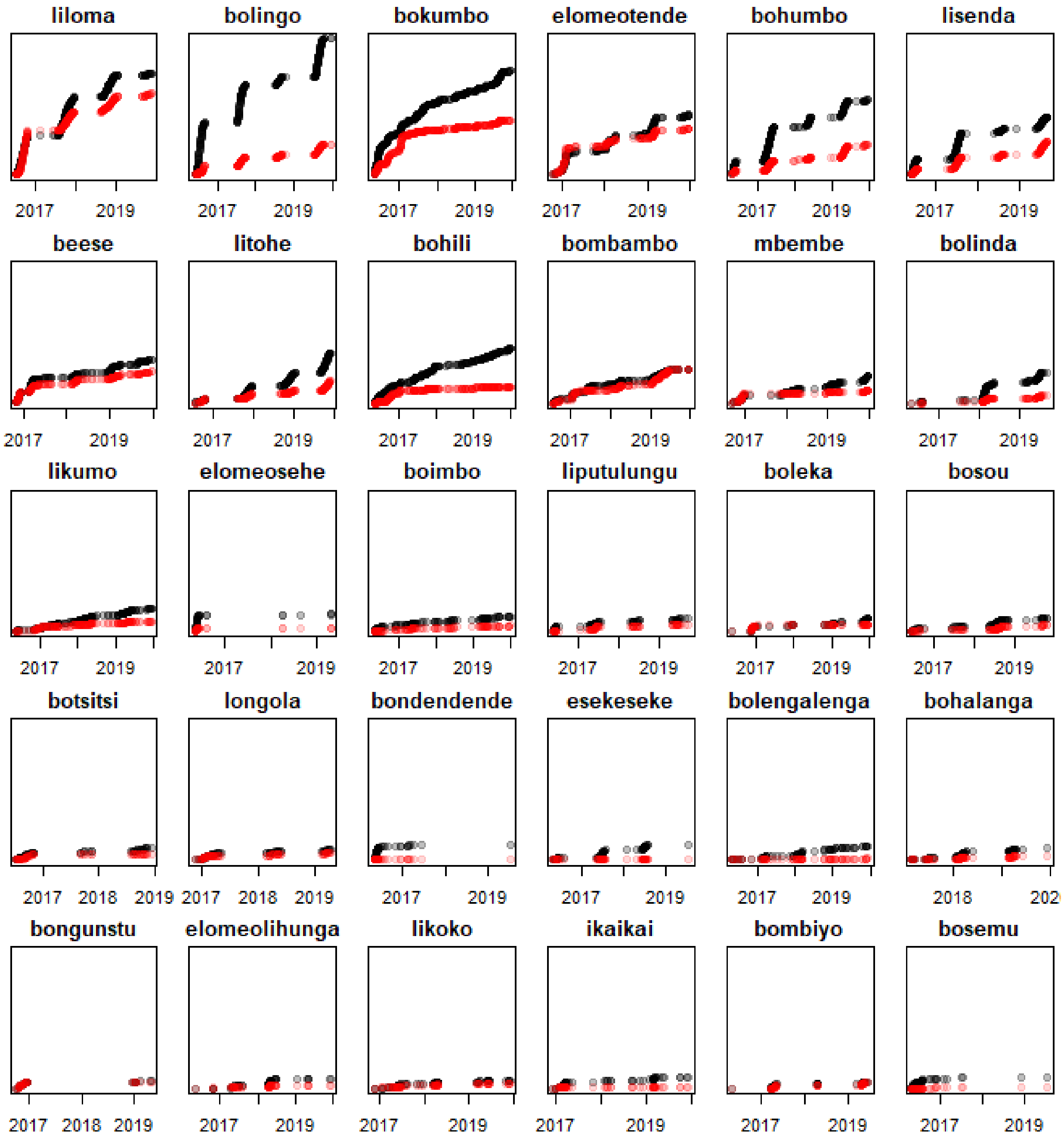

***Fig. S3***. *Accumulation of new cells containing each species over time in the dataset (black) and accumulation of total number of revisits (red) for the top 30 species in the Ekalakala (EKK) diet. Y-axis depicts an absolute scale of number of locations visited (range: 0 to 1070).*

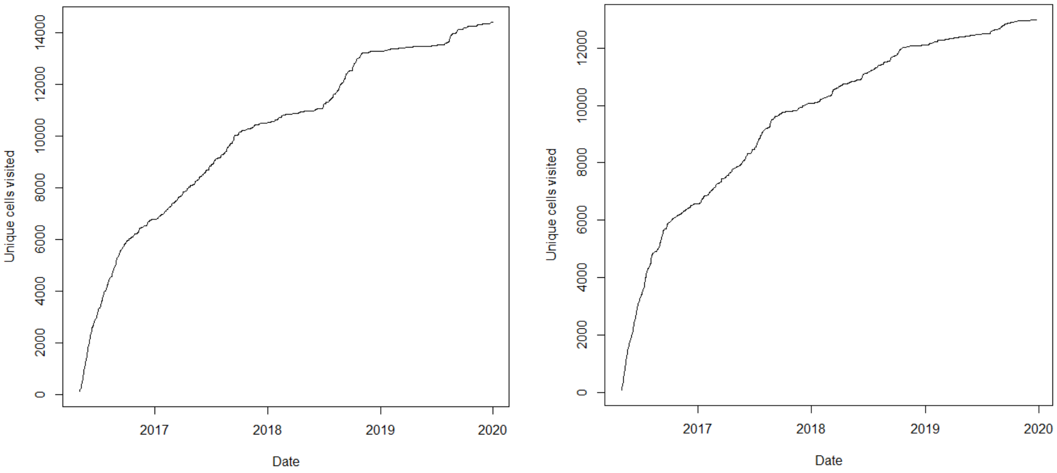

**Fig. S4**. Accumulation of unique cells visited over time for Kokoalongo (left) and Ekalakala (right) social groups.

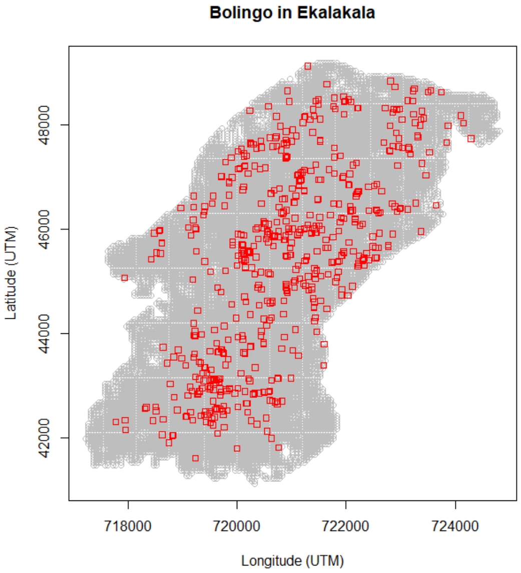

***Fig. S5****. 50 x 50m cells (red squares, not to scale) with 'Bolingo' present relative to the Ekalakala 95% home range kernel (i.e., all visited cells; depicted in gray, also not to scale).*

### 3. Agreement between methodologies

We compared estimates from our vegetation plot dataset with estimates from the CCM dataset on the basis of three different metrics: 1) species densities, 2) measures of dispersion, and 3) distribution of individuals over space. For all subsequent comparative analysis between CCM and vegetation plots, we included only trees and lianas larger or equal in DBH than the smallest visited individual of a given species by bonobos.

#### 3.1 Density

To compare estimated species abundances derived from each dataset (CCM or vegetation plots), we derived three different indices. 1) We used bonobo observational data to create a 'presence index' based on bonobo feeding locations for each food species, estimated as the number of 50x50m cells in which each species was present divided by the total number of cells within the 95% kernel home range of each group (see Figure S5 for an example). 2) We calculated traditional species density estimations using the vegetative plot data as the total number of individuals observed per area surveyed (num. individuals / $km^2$, hereafter "Plot Density"). 3) We calculated the number of 50x50m plots in which each species was present per total number of plots sampled for more direct comparison with the CCM (hereafter "Plot Presence").

To evaluate method agreement, we created pair-wise sets of comparisons of the three density indices by means of Pearson's correlation tests and used the correlation coefficient (r) as a measure of strength of agreement between methods. We conducted the pair-wise comparisons while assessing the influence of sampling effort on method agreement by varying levels of home range usage (kernel % range from 20 until 95 % in increments of 1%) and dietary inclusion (top 10 most consumed species until full diet) for each group. We only considered comparisons with at least 10 species in at least 10 vegetation plots. As density estimates in all three comparisons were right-skewed but occasionally contained cases of true zero or values of one, we added a small constant to the whole dataset (set at half of the minimum non-zero value in the dataset of each method) and log-transformed each set of species densities.

##### 3.1.1 Moving sampling window of density across home range

While the expectation of correlational strength degradation at the periphery of the home range is reasonable, the comparisons of increasing home range inclusion may mask these patterns over the course of the home range as dataset size increases and is averaged over the home range. In other words, if agreement between the two methods degrades at the periphery where sampling effort by the bonobos is expected to be lower, we may have difficulty identifying this degradation pattern if these data are also included with data from the core where sampling effort is highest (and agreement expected to be highest as well). To investigate this possibility, in addition to evaluating agreement across varying levels of home range inclusion, we also compared species density estimates from the two methods using a moving window that ranged along the kernel home range for each group, from 20% to 95% kernel density.

Because peripheral areas naturally encompass larger areas than the core, they are also more likely to encompass a greater number of our systematically placed plots. Therefore, instead of assigning a fixed window size (e.g., including 10% window of comparison, thereby comparing data from 45% - 55% kernel home range, for example), we instead allowed the radius of the sampling window to vary in size to impose uniformity in the number of plots used in each level of the kernel used in the comparison. We allowed the center of this window to range from the 15 to 95% kernel home range of each group, with a varying window size at each interval (1%) to allow for a minimum of 20 plots from which to sample at each interval, with the kernel inclusion radius narrowing once this minimum was met (EKK radius average: 9.5 ± 6.4% (SD), range: 3 - 27; KKL radius average: 9.1 ± 5.5% (SD), range: 3 - 25). Number of plots included in the window per interval averaged 24.3 ± 2.9 (SD; range: 21 – 34) in EKK and 25.4 ± 4.3 (SD; range: 21 - 43) in KKL. Windows were largest in the core of the range where area sampled was most restricted, with the narrowest sampling windows at the periphery where area sampled was largest (e.g., consider the size of the center of a circle relative to the peripheral areas of a circle). We likewise evaluated these datasets across a varying amount of the diet from 10 species to the full diet.

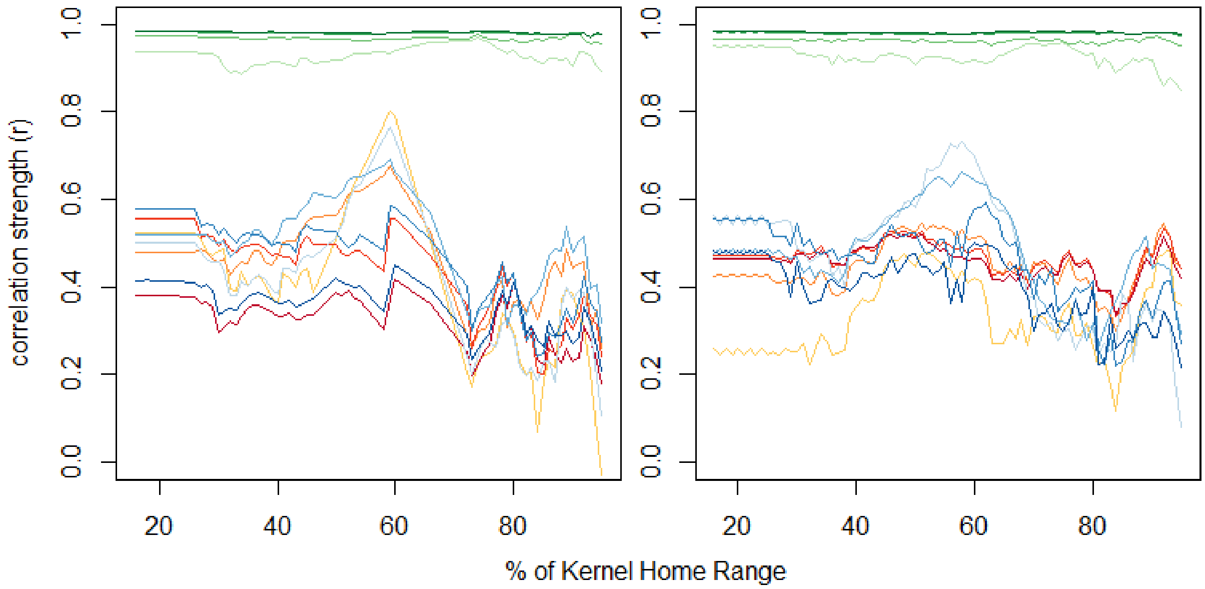

***Fig. S6.*** *Correlation between the three different data types (habitat plot density and p/a proportion, and bonobo p/a) based on variable moving windows (over the percent kernel home range usage. Comparisons are depicted across three methods: CCM and plot density (blue), CCM and plot presence/absence (red), and the control comparison of plot density and plot presence (green). Color gradients (blue to light blue, red to yellow, green to light green) indicate number of species included in home range gradation (ranging from 70 to 15 at approximately 15 species increments).*

We found that once our moving window reached the minimum of 20 plots at ca. 30% kernel home range, correlation strength of the CCM with vegetative plot density estimates increased until they reached a maximum around 60% kernel home range in both groups (Fig. S6). Peripheral areas of the home range were generally lower in agreement than more central areas but did not show persistent decreases with increasing peripheralization in a manner that would suggest consistently poorer sampling in peripheral areas. Sampling agreement was strongest within our moving windows for the most frequently consumed species (e.g., 15 or 30 species) relative to more comprehensive subsets of the two groups' diets (e.g., 55 and 70 species).

3.2 Dispersion

To evaluate the efficacy of the CCM in characterizing dispersion metrics of various food species in the diet, we compared an index of the spatial distribution of species derived from the CCM as well as the same index derived from the vegetation plot data. To do so we used Morisita's index (Morisita 1962). Species in a landscape may be distributed along a gradient from a uniform, over random, to a clumped distribution, and this index uses spatially-explicit samples of each species to quantify patterns of evenness across samples. To allow for standardized and directly comparable sample units to calculate this index between the two data methods, we first aggregated the number of individuals per species visited by the bonobos across three different grid cell sizes (500x500m cells, 1000x1000m cells, and 1500x1500m cells), and calculated Morisita's index using the *dispindmorisita* function of the package 'vegan' (Okansen et al. 2019) for each species with each cell as a sample. Next, we calculated Morisita's index for each species across the three cell sizes using the full vegetation plot dataset within each group's 95% kernel home range and averaged the number of individuals per species per cell to control for the variable number of plots sampled within each cell. We observed an unusual distribution of Morisitia's indices across the three cell sizes, as indices derived from the plot data predominantly ranged between -1 and 1 but several extreme outliers above 1 severely skewed the distribution of the index for this dataset. To allow for more symmetrically distributed values we log transformed all absolute values exceeding values of 1 while retaining the original scale of values between -1 and 1. This allowed a more even normal distribution which we could subject to Pearson's correlations for each species in our dataset.

### 3.3 Distribution

#### 3.3.1 Agreement in species abundances across cells

To evaluate the efficacy of the consumer-centric method to reliably quantify the distribution of food species in a landscape, we applied the identical grid-cell upscaling method as in our dispersion comparison. We compiled the abundance data for both bonobo and vegetation plot datasets in two ways: by aggregating the number of individuals per species per grid cell size (averaged across vegetation plots within each cell) or by marking the presence/absence of a given species per grid cell size. We then fitted a model for each approach (i.e., aggregation or absence/presence) and cell size, using each species as a datapoint. We used the estimated bonobo feeding data abundance per cell as the response predictor and the plot abundance as the test predictor. To account for variation in home range utilization by the bonobos, we controlled for the % kernel home range of each cell by averaging the % kernel value assigned to each of the vegetation plots used to estimate the species abundance within that cell.

We were unable to fit models with identical error structure for all three threshold sizes, and therefore we fitted two types of models to each dataset. Because cells of 500x500m size commonly did not contain individuals of many species utilized by bonobos (and therefore cells for most species contained many zeros) we fitted a zero inflated Poisson model using the function *zeroinfl* of the package 'pscl' (Zeileis et al. 2008;

Jackman 2020), as we found the simple Poisson model to be overdispersed. We found that a zero-inflated Poisson model produced the lowest AIC (corrected for small samples: Burnham et al. 2011) of the model types tested (Poisson, negative-binomial, zero-inflated Poisson, zero-inflated negative-binomial). The count component of the model comprised both predictors (vegetation plot abundance and % kernel), whereas we included only vegetation plot density as a predictor of zero inflation. We assessed model stability by comparing estimates derived from all data with those derived from data excluding areas one at a time. This did not reveal obviously influential cases. We also checked for potential collinearity issues using Variance Inflation Factors (VIF; Quinn & Keough 2002; Field 2005) using the *vif* function of the package 'car' (Fox & Weisberg 2011), applied to a model lacking the zero-inflation component, which did not reveal any issues.

To evaluate agreement between the methods we then used variance explained (Nagelkerke's $R^2$) by the predictor 'abundance from vegetation plots' as a measure of fit between abundance estimated from the bonobo data for each species in each groups' diet. We fitted a null model excluding only vegetation plot abundance in the count component but otherwise identical in structure and took the square-root of Nagelkerke's $R^2$ of the full model (based on comparison with the null model) for comparability with correlation coefficients in our other analyses. To then evaluate the effect of varying 'surveying effort' by the bonobos on the agreement between the two methods, we averaged this correlation strength across species in a varying subset of species, ranging from the top 10 species in each group's diet to the full diet of the group.

As grid cell sizes of 1000x1000m and 1500x1500m frequently contained at least one individual of a species in all grid cells, we could not fit a zero-inflated model; however, we found that a simple model with Poisson error structure was over-dispersed. To resolve these issues, we therefore found that fitting a simple linear model with the response (number of individuals) transformed by a square root produced a model which reasonably conformed to the assumptions of homogeneity and normality of residuals, as well as did not suffer from issues of collinearity. As with the zero-inflated model, we calculated the proportion of variance explained by 'vegetation plot abundance' (r; i.e., square root of the result of the $r^2$ of the full model minus $r^2$ of the null model lacking this term) for each species in the diet of each group and averaged this strength according to dietary inclusion.

#### 3.3.2 Agreement in species presence across cells

To also evaluate agreement between methods on simple presence of a species in a cell, we fitted a generalized linear mixed model with binomial error structure (Baayen 2008) for each grid cell size and each social group. The response in this model was the presence or absence of a species in a given cell as predicted by the bonobo observational data (with one datapoint per species per cell), and presence as measured by vegetation plot and % kernel as test predictors. In these (six total) binomial models we included cell ID and species as random effects and included random slopes for presence/absence in the plots and their correlation within the random effect of species. All six models conformed to assumptions, were suitably stable, and did not suffer from issues of collinearity.

As a last validation of CCM distribution agreement with estimates from vegetation plots, we identified when bonobos missed the presence of a species in a cell that had been identified in habitat plots. To this end, we determined the proportion of missed cells using the number of grid cells where bonobos did not feed on a species determined present by the vegetation plots divided by the total number of cells as a response. We then calculated the proportion of missed species occurrences in cells per species, as well as evaluated potential sources of biases in likelihood to miss a species in a cell (see below).

### 4. Identifying sources of bias

To compare the DBH of food trees and lianas across CCM and vegetation plot data, we fitted a linear mixed model with Gaussian error distribution separately for each social group with the log transformed of the DBH as the response and the method as the test predictor, while controlling for the random effect of species. These models revealed that trees visited by bonobos were significantly larger than trees measured in the plots (EKK: t=-17.71, p<0.001; KKL: t=-20.38, p<0.001), but by only an average of less than 1cm in both groups (Table S1). For 23.1% of consumed species, we found more individuals in the plots that did not reach the minimum size consumed than those who did exceed this minimum threshold.

***Table S1.*** *Correlation of DBH measured in plots with DBH of consumed tree and lianas by bonobos during observation (CCM).*

| | EKK | | | KKL | | |
|---|---|---|---|---|---|---|
| | **Estimate ± SE**** | **t** | **p** | **Estimate ± SE**** | **t** | **p** |
| Intercept | 4.54 ± 0.09 | - | - | 4.57 ± 0.09 | - | - |
| Method (Plot)* | -0.15 ± 0.01 | -17.71 | <0.001 | -0.18 ± 0.01 | -20.38 | <0.001 |

*Reference category: CCM
**Estimates and standard errors are on the original log-transformed scale

4.1 Species characteristics

Because bonobos presumably do not 'sample' available resources equally, characteristics of a species likely influence the patterns in which it is sampled by the bonobos. Species which are highly clumped, exceptionally variable in quality or size, or rare in the landscape can be expected to be sampled differently by a consumer than would be expected from vegetation plot sampling. Therefore, we expected the correlation of density, dispersion, or distribution metrics between our sampling methods to vary depending on food species characteristics. We therefore quantified seven characteristics of each species to evaluate how they contributed to rates of data accumulation and agreement between our sampling methods.

The first of the variables we considered was the *lifeform* of the species (tree or liana), as while tree form allows direct and finite measurement of size, liana species in tropical forests frequently root in multiple locations and extend over large areas of the canopy, thereby complicating our ability to accurately estimate food production, size, and attribute a single location to an individual liana. We therefore expected the correlation between CCM and those based on vegetation plots to be higher in trees.

It may also be possible that the *dispersion* pattern of a species affects how reliably it is sampled by a consumer. We therefore used the transformed values of the Morisita's index assigned to each species from the plot dataset (described above) as a proxy of species dispersion in our models.

We also created a metric which quantified *seasonality* of consumption for each species. The type and seasonality of consumption may have a significant influence on how a species is 'sampled' by bonobos. In a tropical area like the Congo basin seasonal variation is likely not restricted to a single cycle per year but may also show shorter or longer cycles that can differ between species. Hence, to identify seasonal cycle length one must first decide which cycle length to assume for each species. To this end, we first visually inspected the data, and based on that we decided to explore cycle length ranging from 100 to 450 days with an increment of 1 (day). For each combination of species and cycle length we conducted the following steps: i) we converted the dates in which a species was consumed to radians (i.e., an 'angle') by dividing Julian dates by the cycle respective length and then multiplying by 2 x pi, ii) we fitted a model with the sine and cosine of the resulting radians included as predictors (Stolwijk et al. 1999) and iii) we determined Nagelkerke's $R^2$ by comparison with a null model comprising only the intercept. The models were fitted with negative binomial error distribution and logit link function using the function *glmer* of the package 'lme4' (McCullagh & Nelder 1989; Bolker 2008).

Separately for each species we then inspected the $R^2$ plotted against the cycle length. For many species this revealed multiple peaks, but for a majority of species we found a clear peak at a cycle length of ca. 365 days. We hence decided to use this cycle length for all species when quantifying the magnitude of seasonal variation in their consumption. Initially, we considered two ways of quantifying the magnitude of seasonal variation, the $R^2$ as described above and the difference between the maximum and the minimum of the fitted values with respect to the seasonal term (i.e., amplitude). However, the latter was to a considerable extent driven by number of visits to a given species and hence we decided to use the former, with values closer to 1 indicating strongly seasonally consumed species, and values of 0 representing species which are not consumed seasonally.

The consumed plant part may also have an influence on how a species is 'sampled' by bonobos. To account for consumption type we considered whether a species was predominately consumed for its *fruit* (yes/no) or for another plant organ (such as leaves or flowers).

Next, the density of a species in a landscape is likely to predict how extensively a consumer can sample it, with the expectation that if a species is exceptionally abundant it is likely that consumers will under-sample it in the landscape, and the converse may be true for exceptionally rarely consumed species. We therefore used the number of individuals of each species in the vegetation plots divided by the total area of all vegetation plots within the 95% kernel home range of either bonobo community as a measure of the species' *density* in the landscape, log transformed to rectify a severe right-skew in the density distribution.

Additionally, bonobos likely do not randomly select available food items from their landscape but are selective in their foraging decisions based on various characteristics of the species, such as size or nutritional content. Therefore, we used the *coefficient of variation of DBH* (i.e., SD divided by the mean) per species as a proxy of potential selectivity in feeding selection amongst consumed individuals.

Lastly, as species are disproportionately consumed within the diet, we accounted for the *frequency of consumption* by each group by log transforming the number of occasions in the full dataset (2016-2019) that each species was consumed.

These seven variables were used in several models to identify potential biases in the efficacy of using bonobo observational data for ecological sampling. For all subsequent analyses, we restricted these analyses to only those species for which at least 10 separate locations were visited to allow for sufficient data depth in each comparison. First, to evaluate how these seven species characteristics may contribute to the speed at which data accumulate in the CCM dataset, we measured the number of days required for each species to arrive at arbitrary thresholds of the total number of unique locations in the dataset. Because these thresholds are completely arbitrary as well as to ensure our approach is robust against stochastic variation in the shape of the data accumulation pattern, we used multiple thresholds (80-95%, increments of 5%) and describe only broad patterns across these thresholds for the two groups (i.e., patterns which remain consistent across 4 or more of the 8 total models). We fitted generalized linear models with a Gamma error distribution and log link function, and with the number of days to arrive at the threshold of interest as a response, and the test predictors as the seven species characteristics metrics, run separately for each social group (eight models total). We found no issues with collinearity (all VIFs<2), however three data points with exaggerated leverage values were removed from all models; once these data points were removed remaining values indicated no remaining influential cases (Quinn & Keough 2002; Field 2005). Model estimates of some non-significant predictors were found to be moderately unstable, but all statistically significant predictors were sufficiently stable in all models. As all subsequent inference is drawn based upon only averaged estimates across all eight models, we have ignored estimates which were indicated to be unstable when calculating model averages. Lastly, we compared each full model with a respective null model lacking all test predictors (Forstmeier & Schielzeth, 2011) using a likelihood ratio test to test the overall significance of the impact of the seven predictors in each model.

As evaluating the speed at which species arrive at these thresholds may be especially susceptible to patterns of seasonal consumption, we additionally evaluated the number of days a species needed to reach “X”% of locations relative to the date at which that species would have arrived at this same threshold if data accumulated evenly and continuously throughout the duration of the dataset. To this end, we fitted general linear models (i.e., with Gaussian error distribution) with the same model structure as the threshold models described above. We evaluated model validity and stability using various diagnostic tools (Cook's distance, DFBetas, DFFits, leverage and Variance Inflation Factors, distribution of residuals, residuals plotted against fitted values), and verified that models conformed to assumptions. We additionally found that one model (90% threshold, EKK) had one extreme residual outlier and re-fitted the model after excluding this outlier. No subsequent model indicated overly influential cases or deviations from assumptions of normality and homogeneity of residuals (Quinn & Keough 2002, Field 2005), although some non-significant ($p>0.10$) model estimates were unstable in most models.

Species characteristics may influence the likelihood for the two methods to provide similar estimates of density. To therefore evaluate the contribution of species characteristics to the strength of the correlation between the estimated densities of the two methods (vegetation plots and feeding location data), we created a measure of difference between estimates of density deriving from vegetation plots or from the bonobo data by subtracting the CCM density of each species from the density estimated by vegetation plots, each z-transformed (to a mean of zero and a SD of one), and obtaining the absolute value of this difference. We calculated the values of both methods for each species using data from the full home range of each group including the top 70 species in the diet. The reason for z-transforming the two densities before determining their difference was that this removed differences in scale of the CCM and vegetation plot density. The difference between them now represents the difference of density of each species relative to the average of densities per method. We then fitted a general linear model for each social group with the difference measure as the response (log transformed to overcome a strongly right-skewed distribution), with the seven species characteristic as test predictors. For both models, we conducted identical model checks as described above, and found that both models were stable, conformed to assumptions of normality and homogeneity of the residuals, and were free from issues of collinearity.

Species characteristics may likewise affect how likely bonobos are to miss individuals of a species within their home range. Therefore, to evaluate the effects that influence the likelihood of bonobos to miss the presence of a species in a cell (percentage of cells in which presence was missed as response), we also fitted general linear models across three grid cell sizes and both groups with the seven species characteristics as test predictors only for species that occurred in at least 10 cells and were consumed at least 10 times and evaluated model stability and conformity to model assumptions. In four of the six models we found cases of data points with exceptionally high leverage (DFFits), and after removing these cases ($n=1$ or 2 each) found that our models were suitably stable with the exception of some non-significant terms which we do not discuss below.

#### 4.1.1 Results

Full models were found to be significantly different than the respective null model lacking all seven test predictors for predicting data accumulation patterns across species (Table S2). Three characteristics consistently predicted variation in species accumulation speed: landscape abundance, size variability, and food item consumed. Across most conditions, species with higher abundance of individuals in the landscape accumulated slower than those of lower abundance (average estimate: 1.1 ± 1.0 days; range: 1.1 – 1.2). We found that for each unit increase in the coefficient of variation of DBH, data on species accumulated an average of 2.3 ± 1.3 days slower in Ekalakala (range of model estimates: 1.8 – 3.1), indicating that species more variable in size accumulated data more slowly than species more uniform in size, although these results were ubiquitous but unique for Ekalakala only. Further, we detected a significant or trending relationship in five of the eight models for slower accumulation speeds in species consumed for their fruits rather than non-fruit items (estimate average: 1.4 ± 1.1, range: 1.2 – 1.7). No species characteristics were found to explain variation in accumulation speed relative to the speed at which data would have been collected if collected at a consistent rate, as all predictors were non-significant regardless of the threshold or group (full-null model comparisons: range of *p*-values: 0.27-0.94; Table S3).

***Table S2 (next page).*** *Full model results of predictors influencing the number of days until bonobos have visited (a) 80%, (b) 85%, (c) 90%, or (d) 95% of the total number of unique locations for each species. Significant or trending (marginally non-significant) terms are indicated in italicized bold text.*

| | Ekalakala | | | Kokoalongo | | |
|---|---|---|---|---|---|---|
| **(a) 80% threshold** | **Estimate ± SE** | **t value** | **Pr(>\|t\|)** | **Estimate ± SE** | **t value** | **Pr(>\|t\|)** |
| Intercept | 1807.75 ± 2.18 | - | - | 3367.46 ± 2.08 | - | - |
| Species abundance* | ***1.19 ± 1.05*** | ***3.387*** | ***0.001*** | ***1.2 ± 1.05*** | ***3.453*** | ***0.001*** |
| Distribution pattern** | 0.87 ± 1.10 | -1.554 | 0.128 | ***0.79 ± 1.10*** | ***-2.555*** | ***0.015*** |
| Seasonality | 0.72 ± 1.68 | -0.643 | 0.524 | 1.58 ± 1.56 | 1.026 | 0.312 |
| Variability in size (CV) | ***2.57 ± 1.57*** | ***2.109*** | ***0.041*** | 0.85 ± 1.45 | -0.422 | 0.676 |
| Life form: tree*** | 0.8 ± 1.19 | -1.234 | 0.224 | ***0.67 ± 1.16*** | ***-2.653*** | ***0.011*** |
| Frequency of consumption* | 1.02 ± 1.07 | 0.267 | 0.791 | 1.03 ± 1.07 | 0.411 | 0.683 |
| Food part: fruit*** | ***1.68 ± 1.2*** | ***2.906*** | ***0.005*** | ***1.36 ± 1.16*** | ***2.003*** | ***0.052*** |
| full-null model comparisons: | df=7, ch-sq= 4.329, p=0.002 | | | df=7, ch-sq=3.643, p<0.001 | | |
| **(b) 85% threshold** | **Estimate ± SE** | **t value** | **Pr(>\|t\|)** | **Estimate ± SE** | **t value** | **Pr(>\|t\|)** |
| Intercept | 1132.44 ± 2.01 | - | - | 1532.93 ± 1.98 | - | - |
| Species abundance* | ***1.14 ± 1.05*** | ***2.743*** | ***0.009*** | ***1.12 ± 1.05*** | ***2.342*** | ***0.025*** |
| Distribution pattern** | 0.93 ± 1.09 | -0.877 | 0.385 | 0.91 ± 1.09 | -1.101 | 0.278 |
| Seasonality | 0.84 ± 1.59 | -0.38 | 0.705 | 1.28 ± 1.52 | 0.594 | 0.556 |
| Variability in size (CV) | ***3.16 ± 1.49*** | ***2.87*** | ***0.006*** | 1.36 ± 1.41 | 0.886 | 0.381 |
| Life form: tree*** | 0.85 ± 1.17 | -1.068 | 0.292 | 0.79 ± 1.15 | -1.667 | 0.104 |
| Frequency of consumption* | 1.04 ± 1.06 | 0.698 | 0.489 | 1.05 ± 1.06 | 0.859 | 0.396 |
| Food part: fruit*** | ***1.34 ± 1.17*** | ***1.835*** | ***0.074*** | 1.13 ± 1.15 | 0.838 | 0.407 |
| full-null model comparisons: | df=7, ch-sq=2.951, p=0.005 | | | df=7, ch-sq= 1.794, p=0.014 | | |
| **(c) 90% threshold** | **Estimate ± SE** | **t value** | **Pr(>\|t\|)** | **Estimate ± SE** | **t value** | **Pr(>\|t\|)** |
| Intercept | 1451.2 ± 1.78 | - | - | 1297.44 ± 1.87 | - | - |
| Species abundance* | ***1.13 ± 1.04*** | ***3.105*** | ***0.003*** | ***1.11 ± 1.05*** | ***2.393*** | ***0.022*** |
| Distribution pattern** | 0.93 ± 1.07 | -1.004 | 0.321 | 0.98 ± 1.08 | -0.271 | 0.788 |
| Seasonality | 0.69 ± 1.46 | -0.963 | 0.341 | 1.16 ± 1.46 | 0.39 | 0.699 |
| Variability in size (CV) | ***2.04 ± 1.39*** | ***2.148*** | ***0.037*** | 1.62 ± 1.37 | 1.518 | 0.138 |
| Life form: tree*** | 0.91 ± 1.14 | -0.688 | 0.495 | 0.86 ± 1.14 | -1.209 | 0.234 |
| Frequency of consumption* | 1.03 ± 1.05 | 0.655 | 0.516 | 1.04 ± 1.06 | 0.669 | 0.508 |
| Food part: fruit*** | ***1.29 ± 1.14*** | ***1.940*** | ***0.059*** | 1.2 ± 1.14 | 1.412 | 0.166 |
| full-null model comparisons: | df=7, ch-sq= 1.951, p=0.007 | | | df=7, ch-sq= 1.612 p=0.009 | | |
| **(d) 95% threshold** | **Estimate ± SE** | **t value** | **Pr(>\|t\|)** | **Estimate ± SE** | **t value** | **Pr(>\|t\|)** |
| Intercept | 1564.48 ± 1.56 | - | - | 824.69 ± 1.5 | - | - |
| Species abundance* | ***1.11 ± 1.03*** | ***3.588*** | ***< 0.001*** | 1.05 ± 1.03 | 1.54 | 0.132 |
| Distribution pattern** | 0.94 ± 1.05 | -1.189 | 0.241 | 1.05 ± 1.05 | 0.877 | 0.386 |
| Seasonality | 0.65 ± 1.35 | -1.428 | 0.161 | 0.89 ± 1.28 | -0.467 | 0.644 |
| Variability in size (CV) | ***1.77 ± 1.29*** | ***2.215*** | ***0.032*** | 1.13 ± 1.23 | 0.586 | 0.561 |
| Life form: tree*** | 1.01 ± 1.11 | 0.054 | 0.957 | 1.02 ± 1.09 | 0.181 | 0.181 |
| Frequency of consumption* | 1.02 ± 1.04 | 0.633 | 0.530 | 1.06 ± 1.04 | 1.744 | 0.089 |
| Food part: fruit*** | ***1.23 ± 1.11*** | ***2.022*** | ***0.050*** | 1.12 ± 1.09 | 1.36 | 0.182 |
| full-null model comparisons: | df=7, ch-sq= 1.441, p=0.001 | | | df=7, ch-sq= 0.795, p=0.003 | | |

***Table S3****. Full model results of predictors influencing the number of days elapsed until bonobos have visited (a) 80%, (b) 85%, (c) 90%, or (d) 95% of the total number of unique locations for each species relative to the number of days it would have taken to visit the same percentage of unique locations if visits were evenly* *distributed across the duration of the dataset. Significant or trending terms are indicated in italicized bold text.*

| | **Ekalakala** | | | **Kokoalongo** | | |
|---|---|---|---|---|---|---|
| **(a) 80% threshold** | **Estimate ± SE** | **t value** | **Pr(>\|t\|)** | **Estimate ± SE** | **t value** | **Pr(>\|t\|)** |
| Intercept | -106.433 ± 348.334 | - | - | 3.807 ± 0.393 | - | - |
| Species abundance* | 15.627 ± 25.113 | *0.622* | *0.537* | *0.412 ± 0.029* | *1.303* | *0.2001* |
| Distribution pattern** | 2.471 ± 32.445 | *0.076* | *0.940* | -0.051 ± 0.036 | *-0.893* | *0.3773* |
| Variability in size (CV) | 165.119 ± 257.858 | *0.64* | *0.525* | -0.179 ± 0.158 | *2.118* | *0.0406* |
| Life form: tree*** | 138.450 ± 147.378 | *0.939* | *0.353* | 0.022 ± 0.095 | *-0.436* | *0.6654* |
| Seasonality | -46.219 ± 88.705 | *-0.521* | *0.605* | 0.080 ± 0.277 | *-1.796* | *0.0802* |
| Frequency of consumption* | -6.548 ± 34.813 | *-0.188* | *0.852* | *-0.418 ± 0.036* | *-0.818* | *0.4182* |
| Food part: fruit*** | 135.965 ± 94.985 | *1.431* | *0.160* | -0.065 ± 0.104 | *0.953* | *0.3466* |
| full-null model comparisons: | F(7,43):0.751, p-value = 0.630 | | | F(7,39): 1.447, p-value = 0.215 | | |
| **(b) 85% threshold** | **Estimate ± SE** | **t value** | **Pr(>\|t\|)** | **Estimate ± SE** | **t value** | **Pr(>\|t\|)** |
| Intercept | -237.592 ± 284.081 | - | - | 168.021 ± 435.899 | - | - |
| Species abundance* | 8.169 ± 14.597 | *-0.836* | *0.4077* | 23.028 ± 32.249 | *0.714* | *0.479* |
| Distribution pattern** | 19.429 ± 31.135 | *0.560* | *0.5787* | -5.336 ± 36.34 | *-0.147* | *0.884* |
| Variability in size (CV) | 191.312 ± 254.768 | *0.624* | *0.536* | 441.939 ± 286.49 | *1.543* | *0.131* |
| Seasonality | 466.028 ± 239.516 | *0.751* | *0.4569* | 37.267 ± 158.411 | *0.235* | *0.815* |
| Life form: tree*** | -103.082 ± 94.691 | *1.946* | *0.0584* | -98.602 ± 94.644 | *-1.042* | *0.304* |
| Frequency of consumption* | 4.327 ± 30.358 | *-1.089* | *0.2825* | -21.384 ± 44.869 | *-0.477* | *0.636* |
| Food part: fruit*** | 41.377 ± 95.261 | *0.143* | *0.8873* | -29.555 ± 109.213 | *-0.271* | *0.788* |
| full-null model comparisons: | F(7,42): 0.899, p-value = 0.516 | | | F(6,39): 0.566, p-value = 0.779 | | |
| **(c) 90% threshold** | **Estimate ± SE** | **t value** | **Pr(>\|t\|)** | **Estimate ± SE** | **t value** | **Pr(>\|t\|)** |
| Intercept | -18.937 ± 226.442 | - | - | 266.85 ± 417.91 | - | - |
| Species abundance* | 6.81 ± 16.316 | *0.417* | *0.679* | 33.4 ± 30.92 | *1.08* | *0.287* |
| Distribution pattern** | 10.696 ± 21.056 | *0.508* | *0.614* | 13.53 ± 34.84 | *0.388* | *0.7* |
| Variability in size (CV) | 60.365 ± 167.34 | *0.361* | *0.72* | 322.49 ± 274.66 | *1.174* | *0.247* |
| Life form: tree*** | 36.657 ± 95.778 | *0.383* | *0.704* | 55.98 ± 151.87 | *0.369* | *0.714* |
| Seasonality | -12.421 ± 57.749 | *-0.215* | *0.831* | -65.42 ± 90.74 | *-0.721* | *0.475* |
| Frequency of consumption* | -8.588 ± 22.641 | *-0.379* | *0.706* | -40.02 ± 43.02 | *-0.93* | *0.358* |
| Food part: fruit*** | 29.33 ± 61.695 | *0.475* | *0.637* | 16.02 ± 104.71 | *0.153* | *0.879* |
| full-null model comparisons: | F(7,42): 0.143, p-value = 0.994 | | | F(7,39): 0.487, p-value = 0.838 | | |
| **(d) 95% threshold** | **Estimate ± SE** | **t value** | **Pr(>\|t\|)** | **Estimate ± SE** | **t value** | **Pr(>\|t\|)** |
| Intercept | 131.471 ± 156.836 | - | - | 31.573 ± 253.997 | - | - |
| Species abundance* | 10.425 ± 11.307 | *0.922* | *0.3617* | 4.179 ± 18.791 | *0.222* | *0.825* |
| Distribution pattern** | 2.584 ± 14.608 | *0.177* | *0.8604* | 21.434 ± 21.175 | *1.012* | *0.318* |
| Variability in size (CV) | -42.676 ± 116.1 | *-0.368* | *0.715* | 63.732 ± 166.937 | *0.382* | *0.705* |
| Seasonality | 75.014 ± 66.356 | *1.130* | *0.2645* | -31.297 ± 92.306 | *-0.339* | *0.736* |
| Life form: tree*** | 17.63 ± 39.939 | *0.441* | *0.6611* | 5.985 ± 55.149 | *0.109* | *0.914* |
| Frequency of consumption* | -26.586 ± 15.675 | *-1.696* | *0.0971* | -21.163 ± 26.145 | *-0.809* | *0.423* |
| Food part: fruit*** | 1.847 ± 42.767 | *0.043* | *0.9658* | -19.08 ± 63.638 | *-0.3* | *0.766* |
| full-null model comparisons: | F(7,43): 0951, p-value = 0.478 | | | F(6,39): 0.406, p-value = 0.893 | | |

***Table S4**. Factors influencing difference between tree and liana species density estimates derived from CCM or vegetation plots for both social groups. Significant or trending terms are indicated in italicized bold text.*

| | Ekalakala | | | Kokoalongo | | |
|---|---|---|---|---|---|---|
| | **Estimate* ± SE** | **t value** | **Pr(>\|t\|)** | **Estimate* ± SE** | **t value** | **Pr(>\|t\|)** |
| Intercept | 2.523 ± 1.832 | - | - | 1.271 ± 1.193 | - | - |
| *Species abundance** | ***0.613 ± 0.136*** | ***4.499*** | ***<0.001*** | ***0.522 ± 0.088*** | ***5.923*** | ***<0.001*** |
| Distribution pattern** | -0.082 ± 0.168 | -0.490 | 0.627 | -0.037 ± 0.109 | -0.340 | 0.735 |
| *Variability in DBH (CV)* | ***-1.416 ± 0.75*** | ***-1.888*** | ***0.066*** | ***-1.155 ± 0.48*** | ***-2.405*** | ***0.021*** |
| Life form: tree*** | -0.191 ± 0.453 | -0.420 | 0.677 | 0.181 ± 0.288 | 0.627 | 0.534 |
| Seasonality | 1.881 ± 1.317 | 1.428 | 0.161 | ***2.637 ± 0.843*** | ***3.129*** | ***0.003*** |
| Frequency of consumption* | 0.273 ± 0.178 | 1.530 | 0.133 | ***0.350 ± 0.109*** | ***3.194*** | ***0.003*** |
| Food part: fruit*** | -0.269 ± 0.488 | -0.551 | 0.585 | ***-0.567 ± 0.317*** | ***-1.789*** | ***0.081*** |
| full-null model comparisons: | F(7,43):7.374, p-value < 0.001 | | | F(7,43):18.630, p-value < 0.001 | | |

*Log transformed; ** Adjusted transformation of Morisita's index (see description); ***Reference category: Liana or fruit

Full-null model comparisons were significant across all conditions, indicating measurable effects of the species characteristics to predict the likelihood to miss a species in a cell (all p<0.001; Table S6). Increases in species abundance (log scale) correlated with an increase in the likelihood for bonobos to miss the presence of species in a cell regardless of cell size or group by an average of 8 ± 1% ($SE_{mean}$; $estimate_{range}$: 5 – 11%), but were less likely to be missed in a cell if they were more frequently consumed (log-scale; $estimate_{average}$: 10 ± 1% ($SE_{mean}$; $estimate_{range}$: -7 – 14%). In half of the models, we additionally found that species consumed for their fruits were on average 7 ± 4% ($SE_{mean}$; $estimate_{range}$: 6 – 8%) more likely to be missed than species consumed for other plant parts.

**Table S5**. Predictors of presence of a species in a cell as identified by each bonobo group across three cell sizes (500x500m, 1000x1000m, 1500x1500m).

<table>
<tr><th></th><th colspan="3">Ekalakala</th><th colspan="3">Kokoalongo</th></tr>
<tr><td>(a) 500 x 500 cells</td><td>Estimate ± SE</td><td>z value</td><td>Pr(>|z|)</td><td>Estimate ± SE</td><td>z value</td><td>Pr(>|z|)</td></tr>
<tr><td>Intercept</td><td>0.647 ± 0.307</td><td>-</td><td>-</td><td>1.58 ± 0.353</td><td>-</td><td>-</td></tr>
<tr><td>Presence in plots</td><td>0.434 ± 0.128</td><td>3.405</td><td>0.001</td><td>0.322 ± 0.141</td><td>2.276</td><td>0.023</td></tr>
<tr><td>% Kernel density</td><td>-0.037 ± 0.003</td><td>-11.118</td><td><0.001</td><td>-0.054 ± 0.004</td><td>-14.385</td><td><0.001</td></tr>
<tr><td>full-null model comparisons:</td><td colspan="3">Chi-sq=91.269, p-value < 0.001</td><td colspan="3">Chi-sq=130.59, p-value < 0.001</td></tr>
<tr><td>(b) 1000 x 1000 cells</td><td>Estimate ± SE</td><td>z value</td><td>Pr(>|z|)</td><td>Estimate ± SE</td><td>z value</td><td>Pr(>|z|)</td></tr>
<tr><td>Intercept</td><td>2.186 ± 0.503</td><td>-</td><td>-</td><td>2.967 ± 0.609</td><td>-</td><td>-</td></tr>
<tr><td>Presence in plots</td><td>0.565 ± 0.16</td><td>3.544</td><td><0.001</td><td>0.815 ± 0.179</td><td>4.554</td><td><0.001</td></tr>
<tr><td>% Kernel density</td><td>-0.038 ± 0.006</td><td>-6.179</td><td><0.001</td><td>-0.054 ± 0.007</td><td>-7.256</td><td><0.001</td></tr>
<tr><td>full-null model comparisons:</td><td colspan="3">Chi-sq=36.519, p-value < 0.001</td><td colspan="3">Chi-sq=51.69, p-value < 0.001</td></tr>
<tr><td>(c) 1500 x 1500 cells</td><td>Estimate ± SE</td><td>z value</td><td>Pr(>|z|)</td><td>Estimate ± SE</td><td>z value</td><td>Pr(>|z|)</td></tr>
<tr><td>Intercept</td><td>0.502 ± 4.367</td><td>-</td><td>-</td><td>0.905 ± 0.14</td><td>-</td><td>-</td></tr>
<tr><td>Presence in plots</td><td>0.144 ± 3.736</td><td>3.736</td><td><0.001</td><td>0.784 ± 0.142</td><td>5.524</td><td><0.001</td></tr>
<tr><td>% Kernel density</td><td>0.006 ± -6.148</td><td>-6.148</td><td><0.001</td><td>-0.054 ± 0.007</td><td>-7.323</td><td><0.001</td></tr>
<tr><td>full-null model comparisons:</td><td colspan="3">Chi-sq=37.145, p-value < 0.001</td><td colspan="3">Chi-sq=63.596, p-value < 0.001</td></tr>
</table>

*5 Model Checks*

When relevant, we tested the significance of fixed effects in mixed models by comparing the fit of the full model with that of a null model lacking all test predictors while maintaining an identical random effects structure as the full model (Forstmeier & Schielzeth, 2011) using a likelihood ratio test (Dobson & Barnett 2018). In case of models with a Gaussian error structure, we visually inspected qq-plots and residuals plotted against fitted values to confirm conformity of our models to the assumptions of normality and homogeneity of residuals. For all models we ensured they were sufficiently stable by excluding levels of the random effects one at a time and evaluating the range of model estimates relative to that of the full model. No models suffered from issues of collinearity among predictors.

***Table S6***. *Linear model results of factors influencing the likelihood for the CCM to fail to identify the presence of a species identified by the vegetation plots across three cell sizes (500x500m, 1000x1000m, 1500x1500m) and both social groups. Significant or trending terms are indicated in italicized bold text.*

| | **Ekalakala** | | | **Kokoalongo** | | |
|---|---|---|---|---|---|---|
| **(a) 500 x 500 cells** | **Estimate ± SE** | **t value** | **Pr(>\|t\|)** | **Estimate ± SE** | **t value** | **Pr(>\|t\|)** |
| Intercept | 1.399 ± 0.115 | - | - | 1.289 ± 0.129 | - | - |
| Species abundance* | ***0.106 ± 0.009*** | ***12.347*** | ***<0.001*** | ***0.11 ± 0.011*** | ***9.768*** | ***<0.001*** |
| Distribution pattern** | ***-0.021 ± 0.011*** | ***-1.991*** | ***0.053*** | -0.018 ± 0.012 | -1.538 | 0.132 |
| Variability in size (CV) | -0.031 ± 0.047 | -0.649 | 0.520 | 0.034 ± 0.051 | 0.660 | 0.513 |
| Life form: tree*** | -0.034 ± 0.028 | -1.211 | 0.233 | -0.009 ± 0.032 | -0.272 | 0.787 |
| Frequency of consumption* | ***-0.087 ± 0.011*** | ***-7.751*** | ***<0.001*** | ***-0.065 ± 0.011*** | ***-5.696*** | ***<0.001*** |
| Food part: fruit*** | ***0.06 ± 0.031*** | ***1.964*** | ***0.056*** | ***0.082 ± 0.038*** | ***2.161*** | ***0.037*** |
| Seasonality | -0.027 ± 0.083 | -0.328 | 0.745 | -0.096 ± 0.092 | -1.051 | 0.299 |
| full-null model comparisons: | F(7,43):23.16, p-value < 0.001 | | | F(7,41): 16.73, p-value < 0.001 | | |
| **(b) 1000 x 1000 cells** | Estimate ± SE | t value | Pr(>\|t\|) | Estimate ± SE | t value | Pr(>\|t\|) |
| Intercept | 1.331 ± 0.148 | - | - | 1.274 ± 0.145 | - | - |
| Species abundance* | ***0.066 ± 0.011*** | ***5.799*** | ***<0.001*** | ***0.084 ± 0.012*** | ***7.076*** | ***<0.001*** |
| Distribution pattern** | 0.021 ± 0.017 | 1.278 | 0.208 | -0.009 ± 0.016 | -0.554 | 0.583 |
| Variability in size (CV) | -0.083 ± 0.06 | -1.397 | 0.170 | -0.002 ± 0.054 | -0.029 | 0.977 |
| Life form: tree*** | -0.032 ± 0.037 | -0.855 | 0.398 | -0.015 ± 0.033 | -0.443 | 0.660 |
| Frequency of consumption* | ***-0.135 ± 0.014*** | ***-9.591*** | ***<0.001*** | ***-0.102 ± 0.012*** | ***-8.628*** | ***<0.001*** |
| Food part: fruit*** | -0.017 ± 0.039 | -0.419 | 0.677 | ***0.08 ± 0.038*** | ***2.108*** | ***0.041*** |
| Seasonality | 0.134 ± 0.11 | 1.216 | 0.231 | -0.053 ± 0.098 | -0.539 | 0.593 |
| full-null model comparisons: | F(7,41): 17.61, p-value < 0.001 | | | F(7,40): 15.38, p-value < 0.001 | | |
| **(c) 1500 x 1500 cells** | Estimate ± SE | t value | Pr(>\|t\|) | Estimate ± SE | t value | Pr(>\|t\|) |
| Intercept | 1.424 ± 0.144 | - | - | 0.953 ± 0.182 | - | - |
| Species abundance* | ***0.08 ± 0.012*** | ***6.721*** | ***<0.001*** | ***0.048 ± 0.014*** | ***3.548*** | ***<0.001*** |
| Distribution pattern** | 0.021 ± 0.016 | 1.348 | 0.185 | 0.029 ± 0.02 | 1.487 | 0.145 |
| Variability in size (CV) | -0.087 ± 0.054 | -1.598 | 0.118 | -0.009 ± 0.069 | -0.127 | 0.900 |
| Life form: tree*** | -0.031 ± 0.034 | -0.919 | 0.363 | -0.013 ± 0.042 | -0.315 | 0.754 |
| Frequency of consumption* | ***-0.143 ± 0.013*** | ***-11.045*** | ***<0.001*** | ***-0.093 ± 0.015*** | ***-6.013*** | ***<0.001*** |
| Food part: fruit*** | 0.017 ± 0.038 | 0.45 | 0.655 | 0.006 ± 0.045 | 0.139 | 0.890 |
| Seasonality | ***0.316 ± 0.1*** | ***3.174*** | ***0.003*** | -0.004 ± 0.127 | -0.035 | 0.972 |
| full-null model comparisons: | F(7,40): 22.56, p-value < 0.001 | | | F(7,42):8.556, p-value < 0.001 | | |

*Log transformed; ** Adjusted transformation of Morisita's index (see description); ***Reference category: Liana or fruit

*6 Advantages and disadvantages of the CCM method*

***Table S7.*** *Advantages and disadvantages to the use of (a) traditional vegetation plot sampling and (b) the CCM for acquisition of information on food abundance to a consumer.*

| | | |
|---|---|---|
| **(A) Vegetation Plot Sampling** | | |
| | **Advantage** | **Disadvantage** |
| | Provides an objective measure of abundance of all potential tree species in a landscape | Effort may be wasted quantifying tree species that are irrelevant (i.e., ignored) to the consumer |
| | Allows for the quantification of landscape-level characteristics non-specific to the consumer such as overall species richness, total tree density, and total basal area | Choice of method used may inhibit ability for cross-site comparison when different methods are used, may introduce biases or errors towards certain characteristics of measured species |
| | Methodology can be adjusted and tailored to different end goals and to accommodate various characteristics in the environment or survey targets (trees/lianas) | Survey effort may need to be intensive depending on desired outcomes (e.g., if species of interest are rare, or landscape is large, or detailed sub-landscape comparison is needed) |
| | Generally comparable across landscapes and objective (i.e., non-specific) to the landscape rather than particular consumers (e.g., study species) or social units | Is a static measurement - survey area must be resurveyed if changes in the area occur |
| | No 'burn-in' time required: data are immediately useable once minimum sampling is met | Can only approximate distribution of individuals at a scale fixed to the methodology - requires a priori assumptions of relevant scale to a consumer |
| | Does not necessitate direct observation, so consumers do not need to be habituated | Can measure only abundance but cannot provide information on distribution of potential feeding locations or actual availability of resources to a consumer |
| | Is independent of consumer movement, therefore sampling can target areas of interest | |
| | Able to measure dispersion using finite and spatially-explicit samples | |
| **(B) Consumer-Centric Method (CCM)** | | |
| | **Advantage** | **Disadvantage** |
| | For frequently consumed species, could theoretically be capable of providing a census of all relevant individuals of a given species once data are fully saturated | Data are not generalizable beyond the sampled individuals or social group |
| | Provides temporally dynamic monitoring of distribution of visited (i.e. relevant) feeding locations - can reflect changes within the area of interest over time | For now, only appears suitable for quantification of densities and some species' distributions, traditional methods may still be required if other metrics are desired |
| | Provides dynamic monitoring of availability also if behavior of the consumer changes | Information gained is limited only to consumed species |
| | Data reflect true availability of resources rather than abundances (which are blind to patterns of use and temporally varying variability) | Quality of information may be biased towards frequently consumed species |
| | Because they are targeted by the consumer, the CCM may allow for better capture rates of species otherwise rare in the landscape | Requires a 'burn in' period before reliable and stable estimates can be provided and data are of sufficient depth |
| | Tailored directly to social unit (e.g., individual, community) and reflects selection biases inherent to each social unit | Requires direct behavioral observation, can only be used for groups/individuals habituated to direct researcher observation |
| | Dataset improves over time; relevance does not degrade unless significant changes to landscape or diet occur | |
| | Is easily integrated into existing behavioral observation data collection and does not require data collection supplementary to existing observational data collection schemes | |
| | With data collection teams sufficiently trained on botanical identification of all food items, does not require additional research effort of botanists | |
| | Data are collected directly at the scale most relevant to the consumer and are therefore not aggregated to impose *ad hoc* scales of summarization | |
